# A Bayesian inference method to estimate transmission trees with multiple introductions; applied to SARS-CoV-2 in Dutch mink farms

**DOI:** 10.1101/2023.02.07.527429

**Authors:** Bastiaan R. Van der Roest, Martin C.J. Bootsma, Egil A.J. Fischer, Don Klinkenberg, Mirjam E.E. Kretzschmar

## Abstract

Knowledge of who infected whom during an outbreak of an infectious disease is important to determine risk factors for transmission and to design effective control measures. Both whole-genome sequencing of pathogens and epidemiological data provide useful information about the transmission events and underlying processes. Existing models to infer transmission trees usually assume that the pathogen is introduced only once from outside into the population of interest. However, this is not always true. For instance, SARS-CoV-2 is suggested to be introduced multiple times in mink farms in the Netherlands from the SARS-CoV-2 pandemic among humans. Here, we developed a Bayesian inference method combining whole-genome sequencing data and epidemiological data, allowing for multiple introductions of the pathogen in the population. Our method does not a priori split the outbreak into multiple phylogenetic clusters, nor does it break the dependency between the processes of mutation, within-host dynamics, transmission, and observation. We implemented our method as an additional feature in the R-package *phybreak*. On simulated data, our method identifies the number of introductions with high accuracy. Moreover, when a single introduction was simulated, our method produces similar estimates of parameters and transmission trees as the existing package. When applied to data from a SARS-CoV-2 outbreak in Dutch mink farms, the method provides strong evidence for 13 introductions, which is 20 percent of all infected farms. Using the new feature of the *phybreak* package, transmission routes of a more complex class of infectious disease outbreaks can be inferred which will aid infection control in future outbreaks.

## Introduction

Knowledge of who infected whom during an infectious disease outbreak is an important source of information. Characteristics of the outbreak, such as the generation time distribution, are derived from data on these transmission events [31]. Moreover, risk factors for transmission, such as distance between individuals or time lag since infection, can be more accurately quantified, if the infection chain is known. Several methods exist that use data on the time of symptom onset, contacts, or other proximity information, to reconstruct the most likely transmission links between cases [13, 4, 3]. Currently, genetic data is increasingly incorporated into epidemiological inference as an additional source of information to infer individual transmission events, transmission clusters, and even complete transmission trees [9, 12, 22, 27, 29]. The use of both genetic data (i.e., differences in nucleotides between different samples of the pathogen) and epidemiological data (e.g., time of sampling, contacts, and geographic distance) increases the evidence on who infected whom. Moreover, high-risk contacts and superspreaders can be identified when a model is based on both types of data [16, 15]. Therefore, several statistical methods have been developed which take both transmission and evolutionary dynamics of the pathogen into account [5, 23, 30, 10].

Most methods assume a single introduction to the population of interest. However, there are many outbreaks where this assumption does not hold, e.g., *Staphylococcus aureus* or *Pseudomonas aerigunosa* are often introduced multiple times on a hospital ward when infected patients are admitted [25], highly pathogenic avian influenza (HPAI) outbreaks among farms are initiated multiple times by wild birds [28], and Foot and Mouth Disease (FMD) can be introduced multiple times from outside a district [17]. Control measures focusing on transmission between hosts may be less effective if there are also external introductions.

Currently, several methods to infer transmission trees from both genetic and epidemiological data are available. A method designed by Worby et al. [29] allows for multiple introductions, but it only has phenomenological distributions of genetic distances. There is no underlying mechanistic mutation model for the genetic difference within and between transmission trees. The *outbreaker2* package in R [14] also allows for multiple introductions, but there is only a phenomenological distribution of the genetic distances between trees. Moreover, *outbreaker2* assumes mutation at transmission, thereby ignoring within-host evolution of the virus. A method that uses a phylogenetic tree and within-host evolution is *Transphylo* [6], although transmission links are placed on a fixed phylogenetic tree. Both outbreaker2 and Transphylo can deal with unsampled cases within the population, which can be used to link transmission clusters, although this is different than inferring introductions from an exogenous population. To model multiple introductions from an exogenous population, Mollentze et al. [**?**] extended the transmission model of Morelli et al. [20], which simultaneously infers a transmission and phylogenetic tree. Here, the within-host evolution was modeled by the use of a binary tree, making the use of multiple samples per host problematic. Moreover and most importantly, there is no publicly available software to use the method.

To make optimal use of genetic and epidemiological data while allowing for multiple introductions of a pathogen, we propose a method to simultaneously infer introductions and transmissions consistent with an explicit phylogeny describing the genetic history of all samples. This extended version of the method developed by Klinkenberg et al. [18] aims to infer the transmission dynamics of an outbreak, i.e., who infected whom, from both genetic data of the pathogen and epidemiological data, such as the time of sampling and culling. Inference of the transmission tree and the phylogenetic tree is done simultaneously, concerning four processes: genetic diversity (within and between transmission trees), within-host diversity, transmission, and case observation. Samples from posterior distributions of the model parameters are taken, using a Markov-Chain Monte Carlo (MCMC) method. These samples provide information on how likely certain infection times and infectors of hosts are.

To address the possibility of multiple introductions, we relax the assumption of a single index case. We add an artificial host to the set of sampled hosts, which serves as an infector for all index cases (Figure 1). For this artificial host, we introduce the term ‘history host’, referring to the representation of the history of the lineages within the index cases. Using the history host, multiple outbreaks of a pathogen in the same population are merged into a single phylogenetic tree.

**Figure 1.**
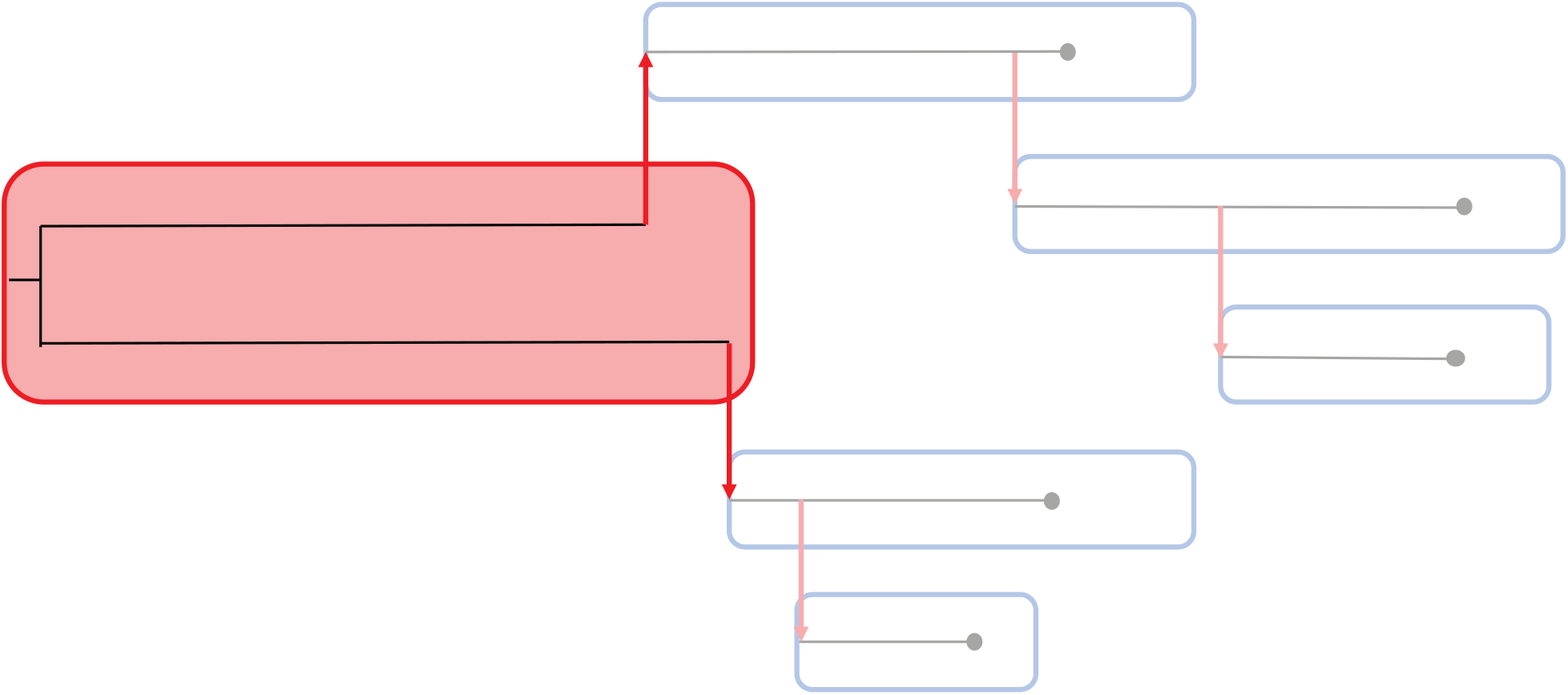
Overview of an outbreak with five sampled hosts and two introductions. The index cases of the sampled hosts (blue squares) are connected via the history host (red square). Coalescence of lineages happens at a different rate in the history host than in the sampled hosts. The black lines give the phylogenetic tree of the outbreak and the red arrows indicate transmissions between hosts.

After evaluation of the performance on simulated outbreaks with single and multiple introductions, we illustrate the application of our method with an analysis of an outbreak of the SARS-CoV-2 virus in the Dutch mink farm industry. From April to November 2020, 63 mink farms tested positive for SARS-CoV-2. To investigate whether the virus was introduced several times into the mink population, we estimated the number of introductions and compared the resulting transmission tree and phylogenetic tree to the phylogenetic tree obtained in [19, 21]. To describe the generation time distribution of infected farms, we used a within-farm model of time since infection, that takes measures to reduce the spread and culling of all animals into account. Furthermore, we implemented the possibility to include multiple sequences per host.

## Results

### Modelling with the history host

To infer the transmission tree of an infectious disease outbreak, we developed a Bayesian method in which four processes define the likelihood of a tree. Mutation events are modeled with a mutation rate *μ*. For the within-host dynamics, we make a distinction between the history host and the sampled hosts. The history host represents either a different population of the same host species, or a different host species (e.g., zoonotic infection), or an environmental source. Therefore, it contains the evolution of the pathogen in the source population, with coalescence happening on a different time scale than within the sampled hosts (see Figure 1). Coalescence, i.e. lineages merging backward in time, is thus described by two rates: rate 1/*r*(*τ*), with *τ* the time since infection, for the coalescence events in the sampled hosts, and rate 1/*r*_history_(*τ*) for the coalescence events in the history host. Timing of transmission is described by a generation time distribution, in the default model a gamma distribution with mean *m_G_* and shape *a_G_*, and for the analysis of the mink farm data we used the generation time described in the methods. Sampling time intervals, as a representation of case observations, are also described by a gamma distribution with mean *m_S_* and shape *a_S_*.

### Improving efficiency of the MCMC

The posterior is sampled by MCMC, with proposals that simultaneously change the phylogenetic and transmission trees. In case there are many introductions, convergence of the MCMC chain to the optimal phylogenetic tree in the history host is usually slow for a random initial configuration of the phylogenetic tree. We solved this issue by (1) initializing the MCMC chain by making each host an introduction and using the neighbor-joining (NJ) tree for the phylogenetic tree in the history host, and (2) implementing the paralleled Metropolis Coupled Monte Carlo Markov Chain (p(MC^3^)) algorithm to give more freedom to the chain [1]. We tested for convergence by comparing the likelihood reached by each algorithm, to the likelihood reached by an MCMC chain starting with the simulated (true) phylogenetic and transmission trees. It turned out that the NJ initialization and the p(MC^3^) algorithm always led to optimal convergence, whereas starting from a random tree and using MCMC sometimes ended up in a local optimum, especially when the number of introductions is high (Table S1). As the tree estimated from the posterior of an MCMC with random initialization did not converge optimally (Figure S1), we say that the configuration of the history host is a bottleneck for performance. Trees may end up in a local optimum of the likelihood. To escape these local optima all following analyses are done with NJ-tree initialization and p(MC^3^).

### Varying number of introductions and coalescent rate

Before assessing in detail the method’s performance to identify the correct introductions and infectors, we compared its performance in relation to different priors. Outbreaks of with 20 hosts were simulated with 5 introductions and a set of default parameters (see materials and methods). The outbreaks were analyzed with uninformative priors on all parameters, informative priors on the mutation rate and mean generation and sampling intervals, and with all parameters set to their true values. Results were compared with respect to identifying the correct infectors, infection times, and parameter values. Only small differences were found between the results of each set of priors for the outbreaks with 5 introductions (Table S2). For instance, the mean numbers of correctly identified infectors were 15, 15, and 15.7, with increasing prior information.

Next, we simulated outbreaks with varying numbers of introductions and varying coalescent rates of the history host. While fixing the number of sampled hosts at 20, we simulated outbreaks with either 1, 2, 5, 10, 15, or 20 introductions. For each number of introductions, we used coalescent rates of 0.004, 0.02, and 0.1 coalescence events per day in the history host, against the background of a mean generation interval of 1 day for transmission events. Thereby we changed the genetic variability of the index cases, by different coalescent rates in the history hosts, resulting in different branch lengths in the phylogenetic tree in the history host. Each combination of a number of introductions and coalescent rate was used for 25 simulated outbreaks, resulting in 450 outbreaks. We analyzed the simulated data with informative priors (Table S2), as in outbreak research most of the time there is some prior information about the generation time and mutation rate.

Analyzing simulated outbreaks with 1 introduction resulted in a mean number (of 25 posterior medians) of 1 introduction, see Figure 2A. This result did not change with the coalescent rate, because there is no coalescence in the history host. With 2 or 5 introductions, the estimated medians were still close to the simulated number. However, with 10 or more introductions the estimated medians were lower than the simulated number of introductions, and a high coalescent rate increased this gap. When all hosts are simulated as an introduction, no more than 40% of all introductions were truly identified by the inference method. This indicates that simulated clusters were merged due to the low genetic variability.

**Figure 2.**
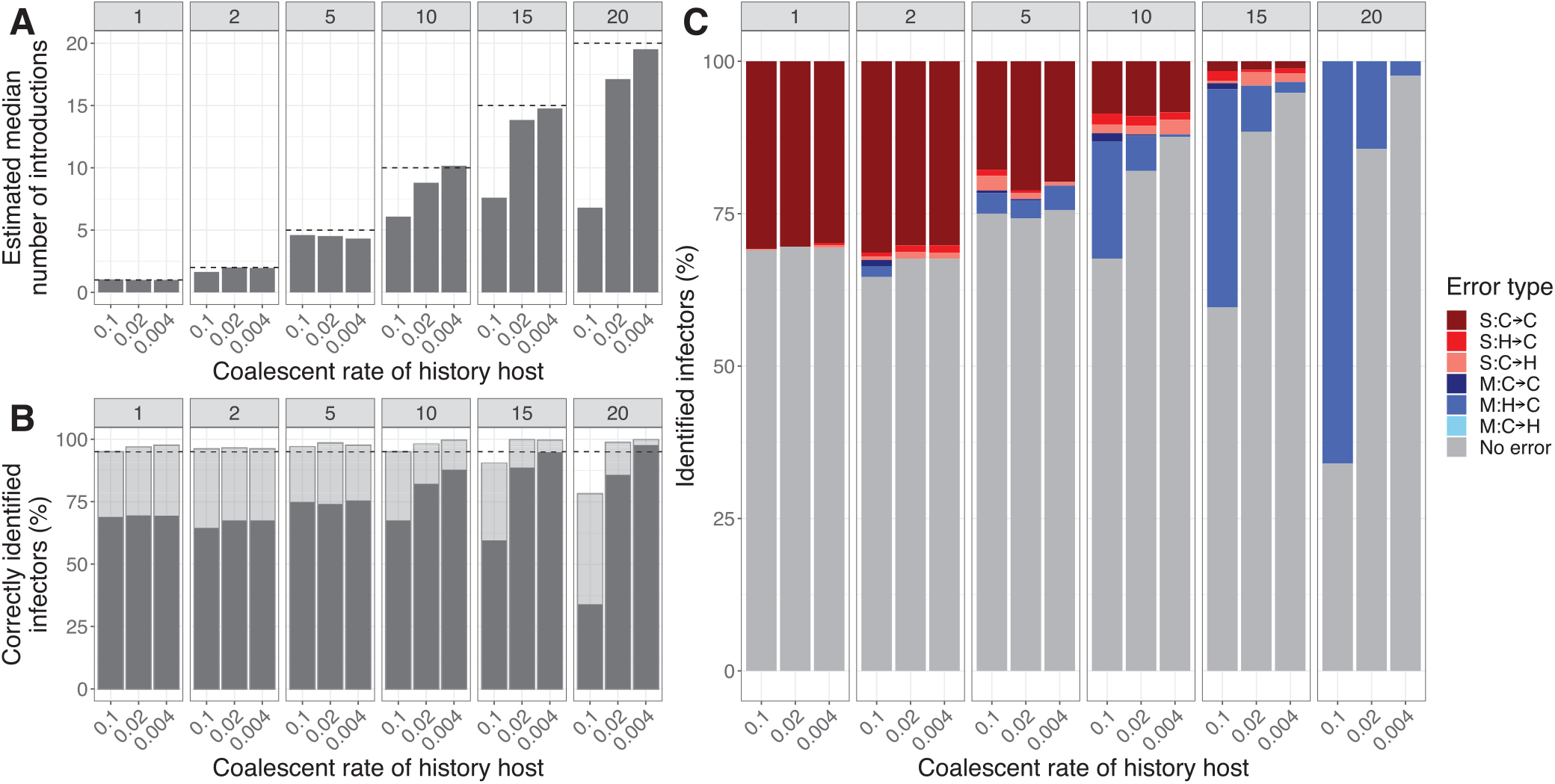
Analysis of simulated outbreaks with a varying number of introductions and coalescent rate (*r_history_*) in the history host. The facets give the results for either 1, 2, 5, 10, 15, or 20 simulated introductions. **(A) The mean estimated median number of introductions.** The black line indicates the simulated number of introductions. **(B) Percentage of correctly identified infectors.** The grey bar indicates cases for which the true infector has the highest posterior weight. The transparent bar indicates cases for which the true infector is contained in the smallest set of candidate infectors with at least 95% of the posterior weight. **(C) Classification of the falsely identified infectors based on the highest support.** The grey bars indicate the correctly identified infectors. S: single transmission cluster involved, M: multiple transmission clusters involved. For the infector of a host: C2C: case becomes case, H2C: history becomes case, C2H: case becomes history.

Approximately 70% of all hosts have correctly identified infectors when there was 1 introduction, and more than 95% of the hosts had their true infectors present in the 95% support set (Figure 2B). This is the set of infectors for a host with cumulative support of at least 95%, with infectors added by decreasing support. For more introductions and low coalescent rates, more infectors were correctly identified, whereas for higher coalescent rates the number of correctly identified infectors decreased.

Several types of incorrectly identified infectors can be distinguished. We define a transmission cluster as the set of hosts derived from one index case. We separate the errors into two classes: involving a single transmission cluster in both the simulated and estimated tree (single, S), or involving multiple transmission clusters in the simulated and/or estimated tree (multiple, M). The simulated or identified infector is then in a different transmission cluster than the case in the simulated or estimated tree. Both classes of error can be subdivided into three subclasses: both simulated and identified infectors are other cases in the data set (case to case, C->C), the simulated infector is the history host and the identified infector is a case (history to case, H->C), and the simulated infector is a case and the identified infector is the history host (case to history, C->H) (see Figure S1). In our analysis, we find that for small numbers of introductions, i.e. 1, 2, and 5, almost all errors are within a single transmission cluster and do not involve an index case (single none). For 10 introductions, this is around half of the errors, while the other half are merges of transmission clusters (multiple simulated). Larger numbers of introductions, i.e. 15 and 20, mostly lead to merged transmission clusters. With the number of introductions approaching the number of sampled hosts, there are only very few transmission events, such that it is hard to estimate the mutation rate or the coalescent rate in the history host correctly. An overestimation of the mutation rate, or an underestimation of the coalescent rate, makes it more likely that index cases are placed in the same cluster, causing merges. Fewer index cases imply more transmission events to estimate the correct parameter values. However, even if all parameters were fixed at their true value, an incorrect infector sometimes has the highest posterior probability (Figure S2).

So, for low numbers of introductions, in these simulations up to 5, the model can reliably infer the number of introductions when informative priors are given for the model parameters. The number of introductions tends to be underestimated if there are many, due to the merging of clusters.

### SARS-CoV-2 in mink farms: analysis of simulated data

In 2020, an outbreak of SARS-CoV-2 occurred among mink farms in the Netherlands. Symptomatic infections in minks first occurred two months after the virus was introduced into the Dutch human population, which suggests that the outbreak was a spillover from humans to mink. To investigate whether there were multiple introductions of the virus into the mink farm population, we applied our extended method to sequence data collected from minks together with their time of sampling. Culling times of the farms were also known. To assess the accuracy of our method on outbreaks with sizes similar to the SARS-CoV-2 outbreak, we simulated and analyzed outbreaks with comparable settings (see material and methods). Again, we tested different numbers of introductions, for which 10 outbreaks each were simulated and analyzed. The results are shown in Table 1. Compared to the percentages of correctly identified infectors for outbreaks with 20 hosts, the model performs equally well for the larger outbreak size of 63 hosts. Around 70-75% of all infectors are correctly identified with the highest support, and the true infector of a host is present in the 95% CI set for at least 95% of all hosts. Only for a high number of introductions (e.g., 20, or 30 introductions), the performance decreases, due to merged clusters, with 5-10% (Figure S3).

**Table 1.** Summary statistics of simulated SARS-CoV-2 outbreaks in mink farms.

|  | Number of simulated introductions |  |  |  |  |  |
| --- | --- | --- | --- | --- | --- | --- |
|  | 1 | 2 | 5 | 10 | 20 | 30 |
| Estimated number of introductions | 1.2 | 2.1 | 4.5 | 7 | 12.7 | 14.3 |
| Correct infectors with highest support | 75% | 75% | 71% | 74% | 74% | 66% |
| Correct infectors in 95% CI | 96% | 97% | 96% | 97% | 97% | 92% |

### SARS-CoV-2 in mink farms: analysis of the Dutch outbreak

During the first and second wave of SARS-CoV-2 infections in the Netherlands (starting in March 2020 and September 2020 respectively), 63 out of a total of 126 mink farms in the Netherlands were sampled positive for the virus. From the end of April 2020 till November 2020, genetic and epidemiological data were collected on these farms, including viral sequences, sampling times, and culling times. A phylogenetic analysis of the viral sequences showed 5 distinct genetic clusters of farms, based on their separation by sequences from human samples [19]. Classification by PANGO lineages [24] showed that each cluster contained one PANGO lineage, with 2 clusters containing the same lineage (Table S?). One farm, NB-EMC-8, contained samples from 2 different clusters and is therefore split into NB-EMC-8a and NB-EMC-8b in our analysis. Whereas the phylogenetic analysis could distinguish five clusters based on human intermediate samples, suggesting five introductions, it could not rule out multiple introductions within each cluster. For an estimate of the number of introductions without the need for intermediate samples from the source population, we analyzed this outbreak with our extended version of phybreak. We set the following priors on the model parameters: *μ_μ_* = 3 · 10^-6^ substitutions per nucleotide per day, *σ_μ_* = 1 · 10^-6^ [2] and the mean *r*_history_ = 20 coalescent events per day with shape equal to 3 (see materials and methods). The mean of the prior introduction rate distribution is 5*/*180, as five genetic clusters were reported within 180 days, with shape equal to 3. Finally, we set the prior mean sampling time *μ_S_* at 10 days, with standard deviation *σ_S_ =* 2, as infection is expected to happen 1-2 weeks before sampling [11].

The method estimated the time of the first coalescent event in the history host on March 4th, 2020 (Table 2). The reduction factor of infectiousness after sampling *L* was estimated at 1, meaning that the method did not find an influence of sampling on infectiousness. We find 13 introductions in the maximum parent credibility tree (see Figure 3), of which 11 have minimal support above 0.5. The median number of introductions in all cycles was 13, with the first and third quartile being 11 and 14 introductions respectively (Figure S7). Six introductions initiated a transmission chain, whereas the other 7 were single cases. By coloring the host labels, we see that the method divided the hosts into subtrees similar to the phylogenetic clusters found by Lu et al. [19]. Two genetic clusters, i.e. cluster B and cluster D, were merged into a single transmission cluster, and with a genetic distance of only 4 nucleotides they belong to the same PANGO lineage. Genetic cluster C is split into two transmission clusters, with NB-EMC-46 as the index case of one of them. NB-EMC-46 was placed in genetic cluster A, but its samples were found to belong to multiple PANGO lineages, including the lineage of genetic cluster C. This indicates that farm NB-EMC-46 is infected multiple times. The large genetic cluster A is separated into multiple transmission clusters, meaning that not all genetically clustered farms are linked by one transmission chain. We find that the single cases which are part of this phylogenetic cluster have common ancestors with cases in the human population (Figure S5). Time of infection and genetic distance made it less likely that the single farms were part of the transmission cluster of farms. In the later stage of the outbreak, there are two larger transmission chains, for which the exact index case is less certain (Figure S8). There is support for the scenario that these transmission clusters are merged into one. In conclusion, by using a phylodynamic model combining the phylogenetic history of the samples with the transmission history between the farms, we were able to distinguish farm-to-farm transmission routes within a group of farms with a common introduction from the human population.

**Table 2.** Summary statistics of SARS-CoV-2 outbreak in mink farms from real data.

| Parameter Inference | median (95% quantile range) of posterior |
| --- | --- |
| $\mu$ | $5.5 \cdot 10^{-6}$ ($4.7 \cdot 10^{-6}$; $6.4 \cdot 10^{-6}$) |
| $m_S$ | 11.9 (10.2; 14.1) |
| $r_{history}$ | 30.5 (17.2; 53.6) |
| $L$ | 1.0 (0.6; 1.5) |
| **Tree inference** |  |
| Number of introductions | 13 (11; 14) |
| Time of first coalescent event in history | -51.7 (-87.4; -27.9) |

**Figure 3.**
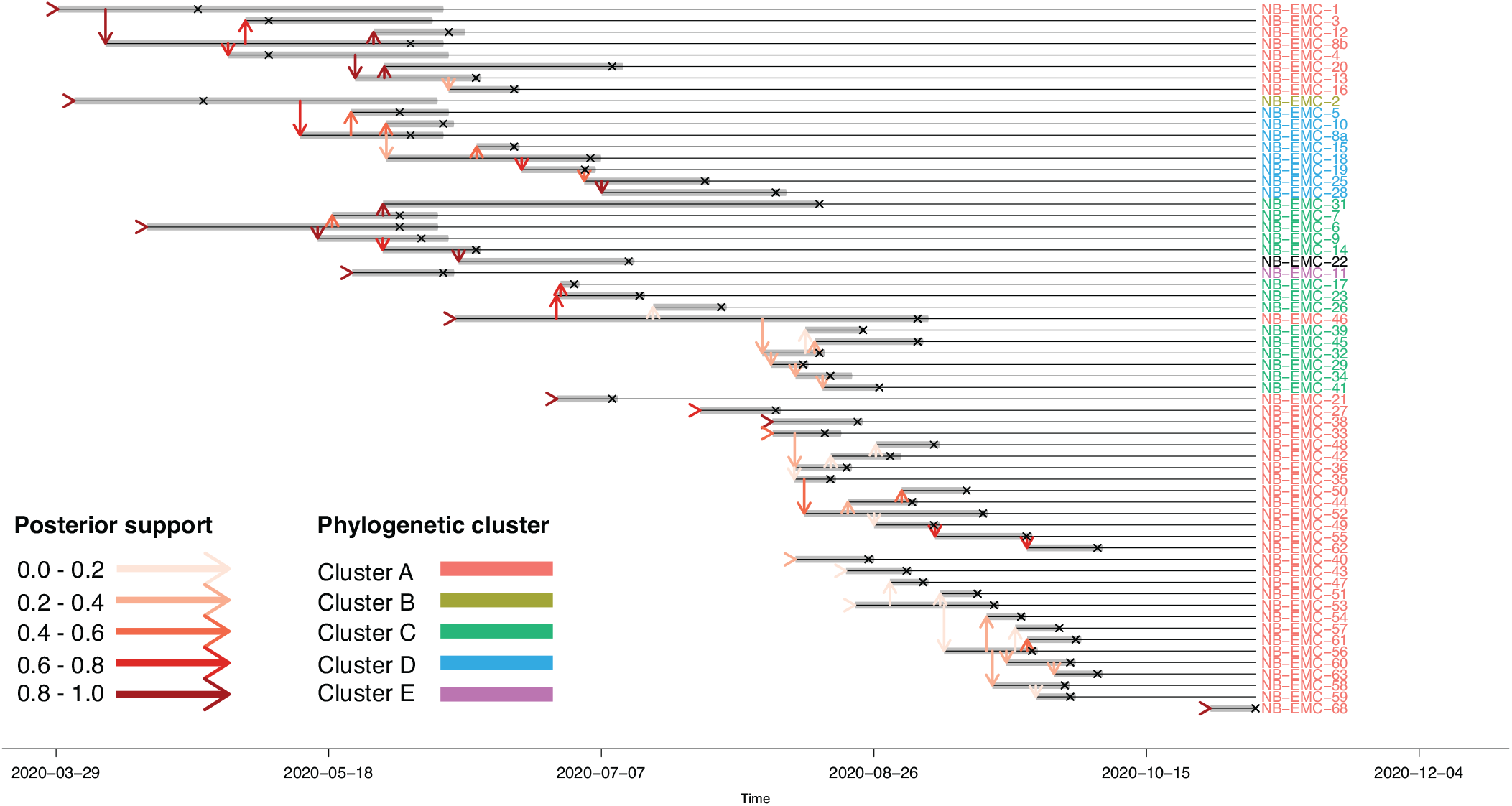
Maximum parent credibility transmission tree of a SARS-CoV-2 outbreak in mink farms. In total 13 introductions are found in the outbreak. Vertical arrows represent transmission links and all arrows are colored according to the support in the posterior distribution. The grey bars show the infectiousness of the hosts and hosts are sampled at the crosses. Host labels are colored according to phylogenetic clusters found by Lu et al. [19].

Our extensions are implemented in the package *phybreak* [18] for the R software [26] and can be found at https://github.com/bastiaanvdroest/phybreak. The package version used, together with the code for the analyses, is found at https://github.com/bastiaanvdroest/phybreak_multiple_introductions.

## Discussion

The method presented enables for the first time to simultaneously estimate the phylogenetic tree and the transmission tree of an outbreak in the case where there may have been multiple introductions. The inference is done without breaking the dependencies between mutations, within-host dynamics, transmission, and observation. By modeling the history of lineages infecting index cases through a phylogenetic tree in a history host, we can distinguish between single and multiple introductions. As an extension to the model of Klinkenberg et al. [18], we now have an easily accessible method for transmission tree inference, with the possibility to assess multiple introductions.

From analyses of simulated outbreaks, we conclude that the model can infer the true number of introductions if there are few introductions compared to the total outbreak size. For an increasing number of introductions, the model increasingly underestimated the number of introductions, but the posterior distribution did include the actual number of introductions. The simulated index cases which were incorrectly identified as non-index cases did have support as an index in the posterior trees. This means that interpretation of the transmission trees should take into account the support as index for cases.

The ability to infer multiple introductions in the analysis of an outbreak is not only useful for finding transmission clusters but also gives valuable information on how to respond to an outbreak. In the case of multiple introductions, measures aimed at reducing transmission events need to be complemented by preventing introduction from outside the target population. Therefore it is of great importance to distinguish between single and multiple introductions of a pathogen in a population. With simulated data sets, we showed that our method is a useful tool to make this distinction: outbreaks with a single introduction are almost always inferred to have a single index case, and outbreaks with multiple introductions are almost never inferred to have a single introduction.

Although the model can distinguish between single or multiple introductions, the accuracy strongly depends on genetic variability. High genetic variability makes it easier to distinguish clusters of hosts, and thus gives more weight to the true number of introductions in the posterior. Low genetic variability, however, will cause sub-trees to be merged and therefore will lead to an underestimation of the number of introductions. As this variability depends on the variation in the external source population, which depends on the mutation rate and effective population size in the history host, it is not possible to state in beforehand how accurate the results will be. When available, strong priors on the mutation rate and coalescent rate in the history host will increase the accuracy, although even with the true values of the model parameters sub-trees will not always be separated. In that case, there is too little information in the genetic and epidemiological data to find all introductions.

Transmission clusters of an infectious disease outbreak in a population are often derived with phylogenetic analyses. However, with closely related index cases, defining clusters may become arbitrary. If obtainable sequences sampled outside of the study population may help to discriminate the clusters by acting as ‘missing links’ between clusters, but discrimination is not so likely if clusters are closely connected. As with the SARS-CoV-2 outbreak in minks, low genetic variability may cause transmission clusters to be merged in the phylogenetic tree, thereby underestimating the number of introductions. We have shown that our method can be used as an alternative approach, which only depends on the genetic data from the study population. Moreover, with the addition of epidemiological data, e.g. sampling times and culling times, it can differentiate genetically similar transmission clusters.

Application of the model to a SARS-CoV-2 outbreak in the Dutch mink farms led to confirmation of previously found phylogenetic clusters, although the phylogenetic clusters are broken down into multiple transmission clusters. These transmission clusters are composed of individual infections along with a larger transmission tree. We split farm NB-EMC-8 based on the genetic clustering of the samples taken on this farm. Without this split, a transmission cluster would have been formed containing multiple PANGO lineages and always having NB-EMC-8 separating the two genetic clusters within that transmission cluster. Farm NB-EMC-46 is also likely to be infected multiple times, as in our results it is the index case of a transmission cluster containing samples from a different genetic cluster than NB-EMC-46. Currently, our method does not allow for multiple infections of a host with different strains, and therefore these clusters could not be separated by the estimation procedure. Extending the method to allow multiple infections of the same host is a challenge for future development. The SARS-CoV-2 outbreak on the mink farms has been studied previously in which samples of humans around and on the farms were used. Here we show that we come to similar conclusions, but do not need samples of the source population to distinguish transmission clusters. Often such data is not available, for example with introductions from other countries, the general population is case of non-notifiable diseases or from wildlife.

The possibility to distinguish multiple introductions of a pathogen into a host population opens up a new avenue for the analysis of outbreaks. However, the method assumes a large population of which a small part gets infected and where contact is equally likely for all pairs of hosts. An outbreak on, for instance, a hospital ward does not meet this assumption with its small population size, in and outflow of patients, and spatial distance between patients. To address these assumptions, the population size has to be accounted for, and contact data, i.e., possible (in)direct contacts between hosts, as well as the geographical location of hosts give a probability of the contact between hosts. Transmission routes can be excluded based on these data sources, such that the certainty of the results increases. In conclusion, we developed a new method for transmission tree inference which makes it possible to estimate the number of introductions of a pathogen during an outbreak. the analysis of the SARS-Cov-2 outbreak in Dutch mink farms shows multiple introductions of the virus, indicating that even with fully controlling farm-to-farm transmission, newly infected farms would arise by new introductions from the human population. Our method opens the way to evaluate outbreaks in such a way that information about new introductions can be derived; knowledge that is useful for policy-making.

## Methods

### Tree inference model

The transmission and phylogenetic tree inference model describes the likelihood of observing an infectious disease outbreak based on the epidemiological and genetic links between hosts and samples. The outbreak dynamics are described by four processes: incidence of new cases by introduction from outside or transmission by existing cases, the observation of the pathogen through sampling, the dynamics of the pathogen within infected hosts and the history host, and genetic mutations in the pathogen. By means of MCMC, we sample from the posterior distribution of parameters and phylogenetic and transmission trees, formed by prior distributions and four likelihood functions for the four processes. The inference is done by a Bayesian analysis, using Markov-Chain Monte Carlo (MCMC) to obtain samples from the posterior distributions of all outbreak parameters and transmission events. We will briefly summarize the likelihood functions, the posterior distributions, and the update steps in the MCMC chain.

Incidence of cases after the first introduction is modeled by two independent processes: additional introduction from outside the study population and transmission between hosts. Additional introductions occur with a rate *λ*_intro_, after the first introduction until the last sample time. We denote by *T* the time between the first introduction and the last sample time, and *k* the number of introductions. Transmission occurs with a dynamic rate, depending on the times since infection of infected hosts, described by the generation time distribution. This is a Gamma distribution with shape *a_G_* and mean *m_G_*. By the use of vector **I** of all infection times, including introductions, and the numeric vector **M** indicating the infectors of all hosts and 0 for introductions, the probability density function of the generation time of a host *i*, with *M_i_* ≠ 0, is *d*_Γ(*a_G_, _mG_*)_(*I_i_* – *I_M_i__*). The likelihood for the transmission tree is therefore:

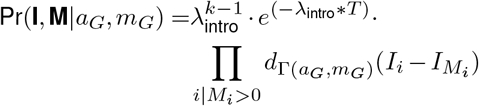

For sampling, we assume that all hosts are detected and sampled at random times after they were infected, according to a Gamma distribution with shape *a_S_* and mean *m_S_*. The likelihood uses the vector *S* of sampling times of all hosts and is therefore:

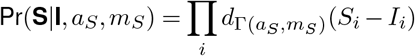

The phylogenetic tree P describes the evolutionary history of all sampled sequences and is built from the phylogenetic mini-trees for each host, connected through the transmission links. The introductions are connected by a phylogenetic tree in a separate ‘history host’. Each mini-tree has tips formed by samples and lineages from secondary cases, and a single root which is a tip in the mini-tree of the infector. Mini-trees are formed by coalescent processes. In (normal) hosts, a rate 1/*w*(*τ,r*) describes coalescence between any pair of lineages within the host going backward in time; in the history host, the rate is constant overtime: *r*_history_. In our analysis, we use *w*(*τ,r*) = *rτ*, the linearly increasing within-host pathogen population size at forward time *τ* since infection of the host. In the phylogenetic tree *P* of the outbreak with the set of nodes *V*, there are three sets of nodes: sampling nodes *V_S_*, i.e. the tips of the tree where sampling took place, coalescent nodes *V_C_* and transmission nodes *V_T_*, where a lineage goes from the infector to its infectee. For node *x, τ_x_* gives the time of the node since infection of the host. The number of lineages in host *i* at time *τ* is then denoted by *L_i_*(*τ*):

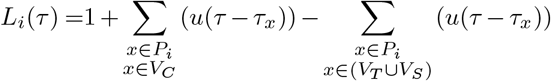

where *u*(*τ*) is the heaviside step function, i.e. *u*(*τ*) = 0 if *τ* < 0, and *u*(*τ*) = 1 if *τ* ≥ 0. The likelihood of each host’s tree is then

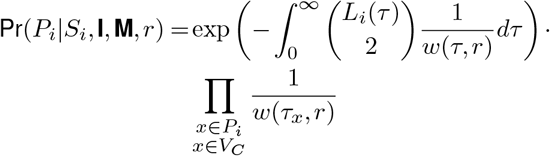

with 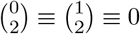. Here, the first term is the probability to have no coalescent event during the intervals in which there are two or more lineages, and the second term is the product of coalescent rates at the coalescent nodes. The prior distribution of the slope *r* is Gamma distributed with shape *a_r_* and rate *b_r_*. Those were set to *a_r_* = *b_r_* = 3 in an uninformative analysis. For the history host, we assume that the coalescent rate is constant over time, so *w*(*τ,r_hist_*) = *r_hist_*. The total likelihood of the within-host dynamics is the product of all hosts’ likelihoods:

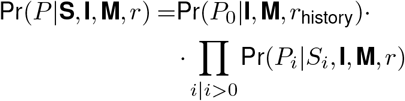

Mutations are described by a Jukes-Cantor model, stating that any of the four nucleotides have equal probability to mutate to, with a fixed mutation rate *μ* for all sites in the set of sequences **G**. For all coalescent and transmission nodes *x*, which occur at time *t_x_* with parent node *v_x_*, the mutation likelihood is:

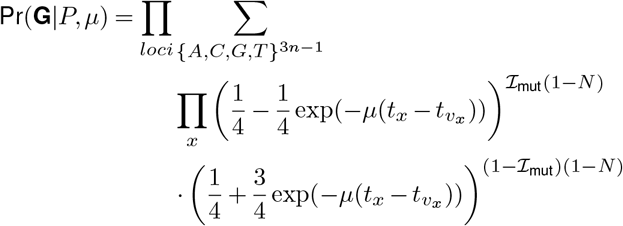

Here, 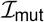 indicates if a mutation occurred on the branch between *x* and *v_x_*, and *N* indicates if a branch ends with a tip with an unknown nucleotide (’n’ in the sequence). We use here a strict molecular clock model, i.e. one mutation rate for all branches of the phylogenetic tree, because on this time scale there won’t any effect of different mutation rates. In the history, changes of mutation rates are met by the coalescent rate of the history host. The likelihood is calculated using Felsenstein’s pruning algorithm [8].

The transmission tree and its parameters are inferred by a Bayesian analysis, using Markov-Chain Monte Carlo (MCMC). From the MCMC we obtain samples from the posterior distributions of the model parameters, the infectors, and the infection times of all hosts. The posterior distribution, with *θ* the set of model parameters, is given by

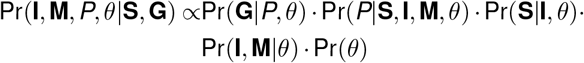

### MCMC sampling

An MCMC chain is run to get the posterior distribution of the model parameters, together with the transmission and phylogenetic tree of the outbreak. The MCMC chains were initialized by first choosing the means of priors for the parameters (except for *μ*), then constructing the transmission and phylogenetic trees, and finally computing a value for *μ*. The trees were constructed by first sampling infection times from the observed sampling times and sample time distribution. All cases were assumed to be index cases (other options are possible within the package), and the topology of the phylogenetic tree was made with the neighbor-joining algorithm using the first sequence of each host. The times of the coalescent nodes were simulated with the coalescent model. This guaranteed an optimized tree topology in the history host, not needing to be reached by sampling in the MCMC chain. The parameter *μ* was for the initial state set to be the tree parsimony (the number of mutations on the tree) divided by the sum of all branch lengths and the genome size. The default prior distributions for the model parameters are found in Table 3. The priors for *m_G_* and *m_S_* are translated into a prior for the rate parameter in the Gamma distribution. More detail about the prior and posterior distributions is included in the supplementary material. Per iteration cycle, each host is picked once in random order as the focal host. A new infection time 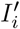 is proposed for focal host *i* and consecutive steps are made according to this new infection time. At the start of a proposal, there are two main ways of updating: within a sub-tree, by following all hosts with a common index case along their transmission links, or between sub-trees. Here we will describe the proposal step for updating between sub-trees, as this is the step where the number of introductions can be altered. The update steps within a sub-tree are as in the original *phybreak* package and can be found in the supplementary information.

**Table 3.**
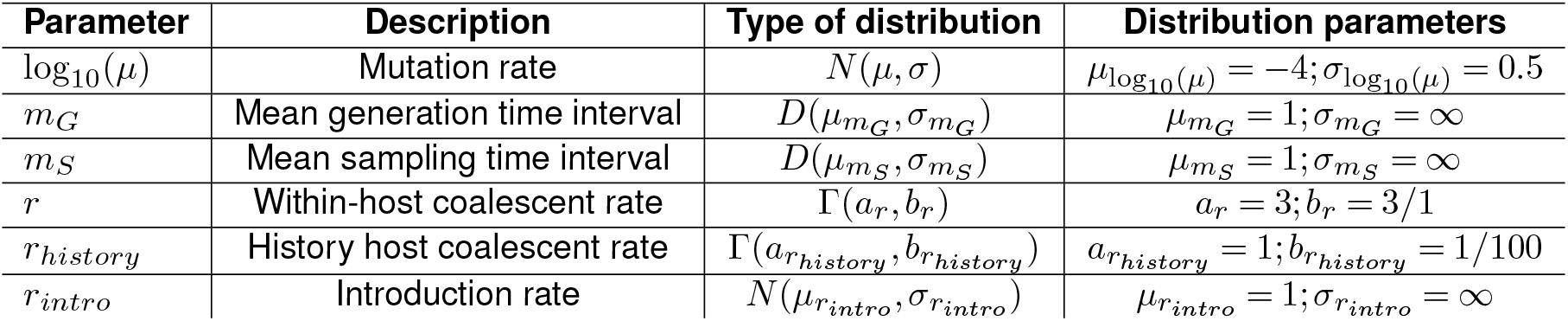
Prior distributions of the model parameters.

| Parameter | Description | Type of distribution | Distribution parameters |
| --- | --- | --- | --- |
| $\log_{10}(\mu)$ | Mutation rate | $N(\mu, \sigma)$ | $\mu_{\log_{10}(\mu)} = -4; \sigma_{\log_{10}(\mu)} = 0.5$ |
| $m_G$ | Mean generation time interval | $D(\mu_{m_G}, \sigma_{m_G})$ | $\mu_{m_G} = 1; \sigma_{m_G} = \infty$ |
| $m_S$ | Mean sampling time interval | $D(\mu_{m_S}, \sigma_{m_S})$ | $\mu_{m_S} = 1; \sigma_{m_S} = \infty$ |
| $r$ | Within-host coalescent rate | $\Gamma(a_r, b_r)$ | $a_r = 3; b_r = 3/1$ |
| $r_{history}$ | History host coalescent rate | $\Gamma(a_{r_{history}}, b_{r_{history}})$ | $a_{r_{history}} = 1; b_{r_{history}} = 1/100$ |
| $r_{intro}$ | Introduction rate | $N(\mu_{r_{intro}}, \sigma_{r_{intro}})$ | $\mu_{r_{intro}} = 1; \sigma_{r_{intro}} = \infty$ |

Three situations describe the possibility to update the transmission tree between sub-trees (see figure 4):

1. The focal host *i* is the history host. In this case, new coalescent times are proposed. Optionally, a new phylogenetic mini-tree can be proposed.
2. The focal host *i* is an index case. An infection time 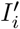 is proposed. If this 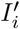 is before the first transmission from host *i*, a new infector 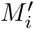 is proposed out of the hosts which are infectious at time 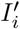. Two situations are now possible:

a. If 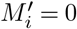, then host *i* remains an index case, with infection time 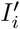.
b. If 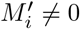, then host *i* is no longer an index case, and there is one introduction less. Host *i* and its descendants will be merged as a branch to another sub-tree.
3. The focal host *i* is not an index case. An infection time 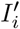 is proposed. If this 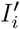 is before the first transmission from host *i*, a new infector 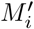 is proposed out of the hosts which are infectious at time 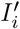. Two situations are now possible:

a. If 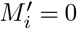, then host *i* will become an index case, and there is one extra introduction. The new sub-tree consists of host *i* and all of its descendants.
b. If 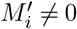, then host *i* either switch to another branch in its sub-tree or switch to another sub-tree. There is no change in the number of introductions.

**Figure 4.**
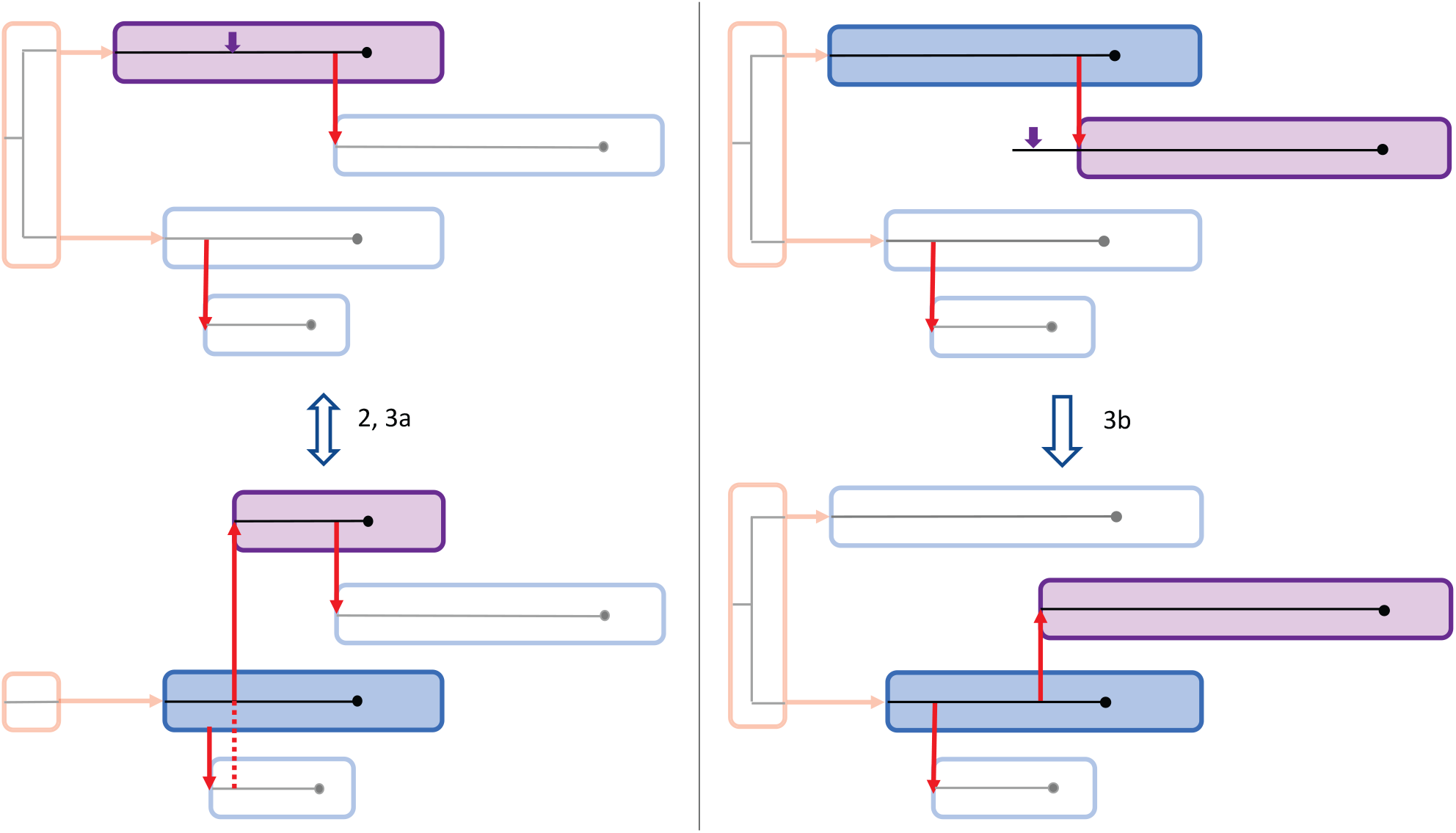
Proposal steps for updates between sub-trees. In purple is the focal host, with the purple arrow indicating the proposed infection time 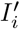. The red arrows indicate the transmission events and the history host is colored red, with the introductions as transmission from the history host. 2: Losing an introduction by proposing a new infector *M_i_* ≠ 0 for an index case. 3a: The reverse of 2, by proposing a new infector *M_i_* = 0 for a non-index case. 3b: Switching sub-trees by proposing a new infector *M_i_* ≠ 0 on a different sub-tree. Situation 3b is also possible within the same sub-tree.

Each proposal step is followed by proposing new phylogenetic mini-trees for all hosts involved. The proposal distributions and acceptance probabilities of all steps are described in the supplemental materials. The MCMC chain is run according to the (MC)^3^ algorithm described by Altekar et al.[1] to improve convergence to the global likelihood optimum. The chains consisted of 35,000 cycles of which the first 10,000 were used as burn-in.

### Construction and analysis of simulated outbreaks

To verify the implementation of multiple introductions in the model, we simulated outbreaks including one or more index cases, and analyzed them by running MCMC chains. The simulation of an outbreak starts with the simulation of a transmission tree:

1. Set an observation size, i.e. the number of hosts, the number of introductions *k*, and the duration of the outbreak *T*.
2. Calculate the optimal population size in which to simulate the outbreak from parameter *R0* and the observation size.
3. Sample *k* – 1 introduction times from the exponential waiting time distribution with rate *λ*_intro_. The introduction time of the first index case will be 0, and other introductions are at cumulative waiting times from the first index.
4. For the index cases, sample the number of secondary cases from a Poisson distribution with parameter *R*_0_.
5. The generation time between two hosts is Gamma distributed with shape *a_G_* and mean *m_G_*. After infection, the sampling of a host takes place after a Gamma distributed time with shape *a_S_* and mean *m_S_*.
6. Repeat steps 3 and 4 for the complete population size, where the infection time for a host is not after *T*. Remove non-index cases without any links.
7. Repeat 3-6 till the desired observation size was given.
8. Add the history host and connect the index cases to this host.

After the simulation of the transmission tree, the phylogenetic tree is constructed by simulating phylogenetic mini-trees for each host. Coalescent times are sampled according to the given coalescent rate 1/*w*(*τ,r*). Edges between sample, coalescent, and transmission nodes are made backward in time. In the history host, coalescence events occur with a constant rate 1/*r*_history_.

For the sequences, we sample the number of mutations from a Poisson distribution with parameter equal to *λ* = *μ* · sequence length · total length of all edges, where *μ* is the mutation rate. The mutations are distributed over the edges, with weights the lengths of the edges. For each mutation, a uniform random locus is changed to a uniform random nucleotide.

We simulated outbreaks with a basic set of parameter values, the same as in Klinkenberg et al. [18], (*mG* =1, *aG* = 10, *mS* =1, *aS* = 10, *R*_0_ = 1.5, *r* = 1, a sequence length of 10^4^ nucleotides and a mutation rate of *μ* = 10^-4^), with new parameters at *λ*_intro_ = 1. The number of introductions varied between the simulations to assess the performance of the model. MCMC chains were run following the (*MC*)^3^ algorithm, with 3 parallel chains with heats 1, 0.5, and 0.333. The chains are 35,000 cycles long, of which the first 10,000 cycles are used as burn-in. Posterior distributions for infectors, infection times, and model parameters are collected from the remaining 25,000 cycles.

### Analysis of SARS-CoV-2 outbreak in Dutch mink farms

As an application of the method, we analyzed the SARS-CoV-2 outbreak in the Dutch mink industry in 2020 [19]. We collected the full viral genomes in minks at 63 farms from GISAID (gisaid.org) and aligned them with MUSCLE [7]. The alignment contains 326 sequences of 29,775 nucleotides long. All positions with N in all 326 sequences are removed because we do not know if there is a mutation at such a position. This left us with 326 sequences of 16,289 nucleotides long. Each farm is sampled at least once, and we have an average number of 5 samples per farm, each farm sampled on a single day. Besides the date of sampling, we also have the date of culling, which is between 1 day and 45 days after sampling, with an average of 4 days. The first 5 farms found to be infected had more than 30 days between sampling and culling, but for the rest of the farms, this was no more than 10 days.

We described the outbreak among mink farms by taking the farms as hosts. The prior distributions of the model parameters are set as follows: we set the mean sampling time interval *m_S_* = 10 days (with a shape *a_S_* = 3), as the time between infection and detection was estimated to be 1-2 weeks [11]. We set the mean introduction rate to 5/180 (with a shape of 3), as five different clusters were found during the outbreak, which lasted for approximately 180 days, by Lu et al. [19]. The coalescent rate parameter *r*_history_ was set to 20. With an expected number of 5 introductions, this rate represented the introduction of the virus in the Netherlands two months before the first positive mink sample. The other prior distributions were set to default.

As the hosts are farms here, we introduced an infectiousness function describing the growth and circulation of the virus within the mink population of a farm. This function replaced the gamma distribution for the likelihood that one farm infected another. We assumed that infectiousness follows a logistic curve, with a reduced level after detection at time *T_s_*, and exponential decline after culling at time *T_c_*:

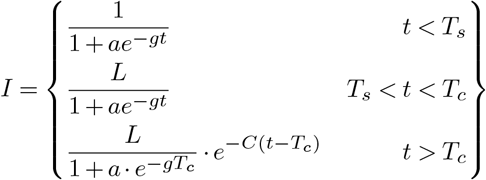

Here, *a* = 1 · 10^-4^ is the initial part of the mink population at a farm being infected, *g* is the growth rate, and *t* is the time after infection of the farm. Parameter *L* is estimated to see if there was some reduction of infectiousness after detection, and *C* is a fixed value. Because the values for *T_s_* and *T_c_* differ per farm, the infectiousness curves differ between the farms. Therefore we normalize the curves, such that the mean AUC of all curves is 1. Then, on average a farm has a distribution of infectiousness that adds up to 1, just as in the default *phybreak* model, while accounting for higher total infectivity of longer infected farms. Another addition used for the mink farms was to include multiple samples per farm. Phylogenetic mini-trees are then built with multiple lineages within a farm, increasing the amount of genetic data. For the sampling time distribution, only the first sample of each host is used.

To test the new model, with a similar history host, and sampling time distribution, we simulated outbreaks with the same parameters as before but with the new infectiousness curve. Culling times were set 15 days after infection, such that the hosts have a fixed infectiousness curve. As for the outbreak size, we used 63 hosts with 1 sample per host. Prior distributions were set with the same parameter values as the analysis of the real data. We set *C* to 5, such that in 5 days after culling the infectiousness of a farm was 0. We varied the number of introductions, from 1, 2, 5, 10, 20, up to 30 introductions. Results of the SARS-CoV-2 outbreak were obtained by running three parallel chains, with 25,000 cycles each, according to the (*MC*)^3^ algorithm. The maximum parent credibility tree is used for visualization, computation of the number of introductions, and comparison to the phylogenetics [19].

## Acknowledgments

This work was performed as part of the research program of the Netherlands Centre for One Health (www.ncoh.nl). We thank Bas Oude Munnik, Francisca Velkers and the ‘One Health mink outbreak investigation consortium’ for providing a prepublication of the mink data.

## Supplementary Information

**Figure S1.**
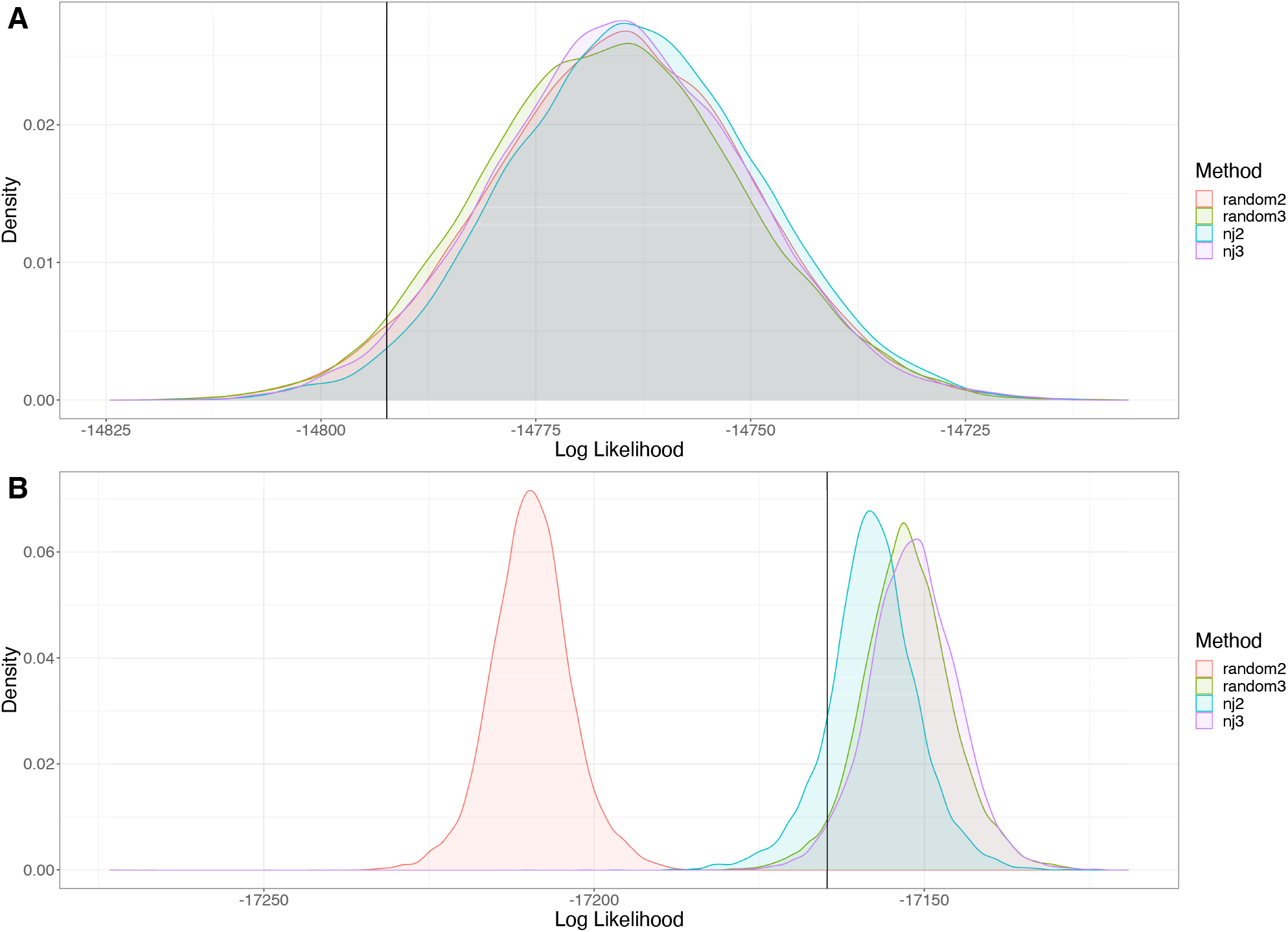
Comparison of MCMC and p(MC^3^) with and without the neighbour-joining tree initialization step. A: For low numbers of introductions (5 of the 20 hosts), there is no difference between methods in the posterior log-likelihood distribution. B: Higher numbers of introductions (15 of the 20 hosts), performance of MCMC with a random tree as initialization of the history host is inferior to either p(MC^3^), neighbour-joining tree initialization of the history host or the combination of both. The latter gives the highest likelihood distribution and is chosen as default option in all analyses. ‘random’ is random tree initialization, ‘nj’ is neighbour-joining tree initialization, ’2’ is MCMC and ’3’ is p(MC^3^).

**Figure S2:**
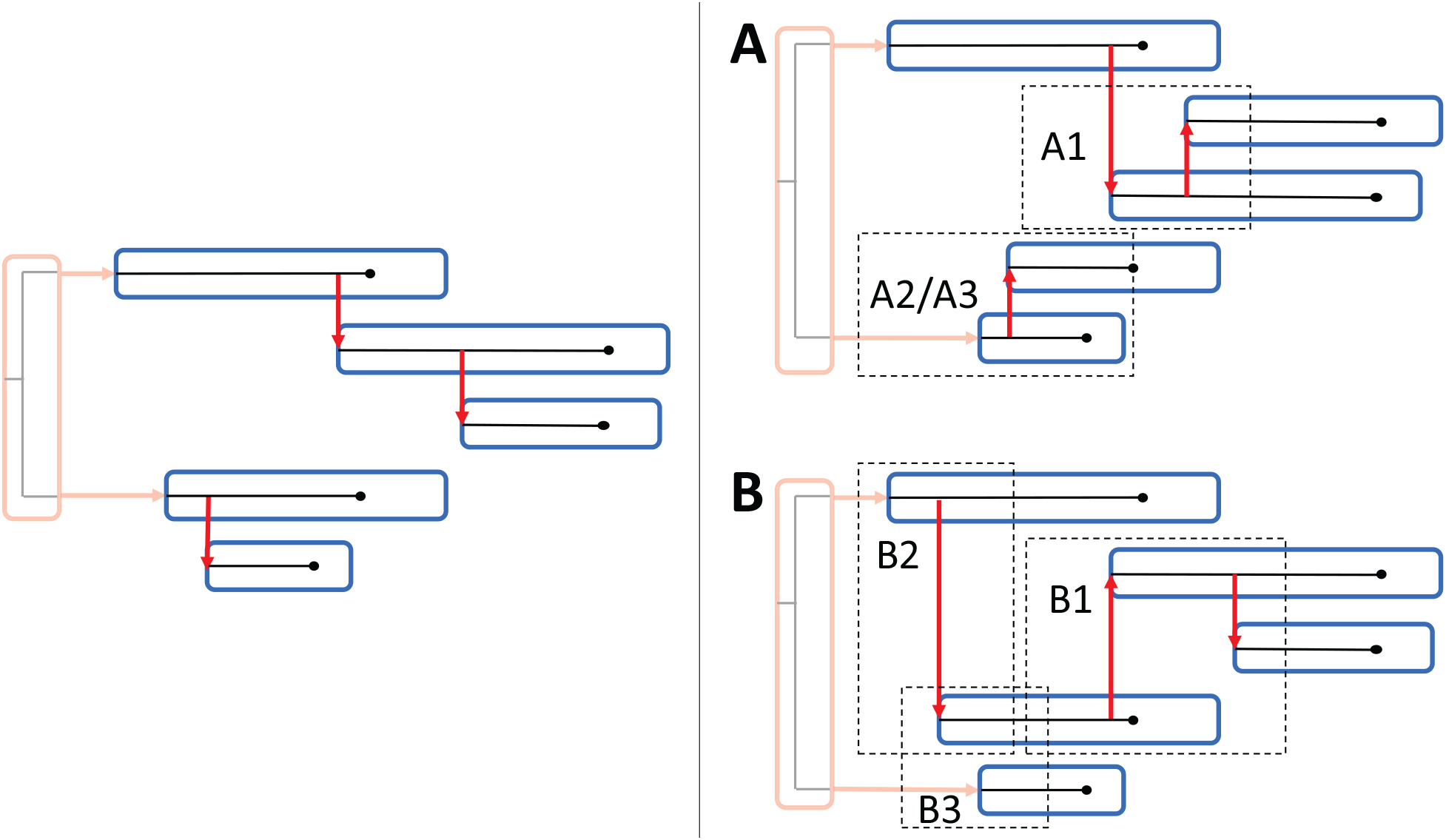
Type of errors in the estimated transmission tree. The left figure represents the transmission tree of a simulated outbreak with 5 cases; there are 2 introductions (clusters) and 3 transmission events. The right figures represents possible estimates of the transmission tree of the simulated outbreak. The vertical ordering of cases in the left and the right figures is identical. The upper right figure shows errors in which an incorrect infector is identified, but the incorrect infector belongs to the same cluster as the true infector (type A errors), the lower right figure represents incorrect identifications of the infector in which the incorrect infector belongs to a different cluster as the true infector (type B errors). In Type 1 errors neither the true infector nor the incorrect identified infector is an index case. For type 2 errors, the host is an index case in the simulated outbreak but not in the estimated outbreak. For type 3 errors, the host is not an index case in the simulated outbreak but is an index case in the estimated outbreak.

**Figure S3:**
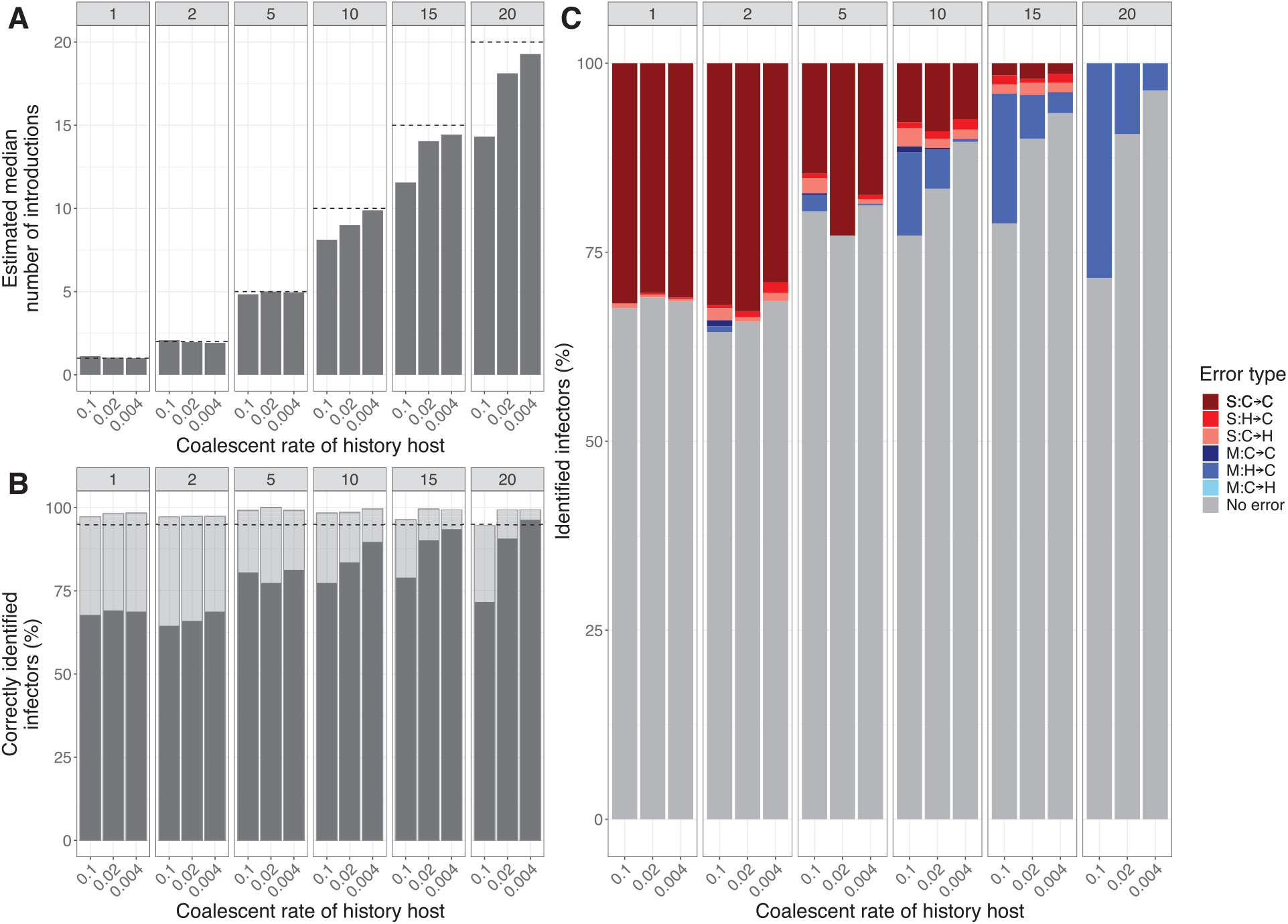
Analysis of simulated outbreaks with varying number of introductions and coalescent rate in the history host. The model parameters are fixed at the simulation values. (A) The mean estimated median number of introductions. The black line indicates the simulated number of introductions. (B) Percentage of correctly identified infectors. The grey bar indicates cases for which the true infector has the highest posterior weight. The transparent bar indicates cases for which the true infector is contained in the smallest set of candidate infectors with at least 95% of the posterior weight. (C) Classification of the incorrectly identified infectors in the maximum credibility tree. The grey bars indicate the correctly identified infectors. S: single transmission cluster involved, M: multiple transmission clusters involved. C->C: simulated and inferred infectors are cases, H->C: simulated infector was history host, inferred infector is case, C->H: simulated infector was case, inferred infector is history host.

**Figure S4:**
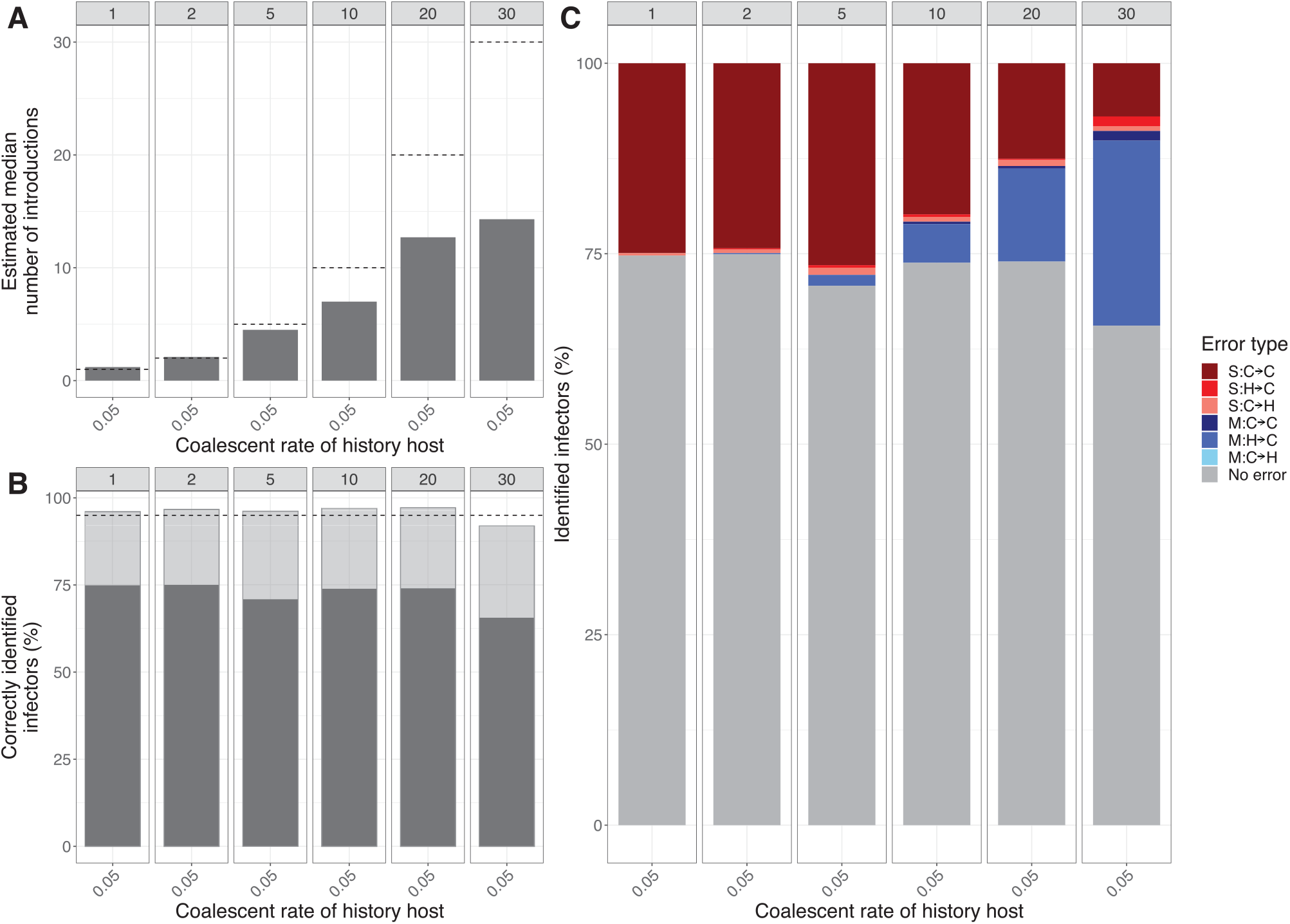
Analysis of simulated outbreaks with similar parameter values as the SARS-CoV-2 outbreak in mink farms. (A) The mean estimated median number of introductions. The black line indicates the simulated number of introductions. (B) Percentage of correctly identified infectors. The grey bar indicates cases for which the true infector has the highest posterior weight. The transparent bar indicates cases for which the true infector is contained in the smallest set of candidate infectors with at least 95% of the posterior weight. (C) Classification of the falsely identified infectors based on highest support. (C) Classification of the falsely identified infectors based on highest support. The grey bars indicate the correctly identified infectors. S: single transmission cluster involved, M: multiple transmission clusters involved. For the infector of a host: C->C: case becomes case, H->C: history becomes case, C->H: case becomes history.

**Figure S5:**
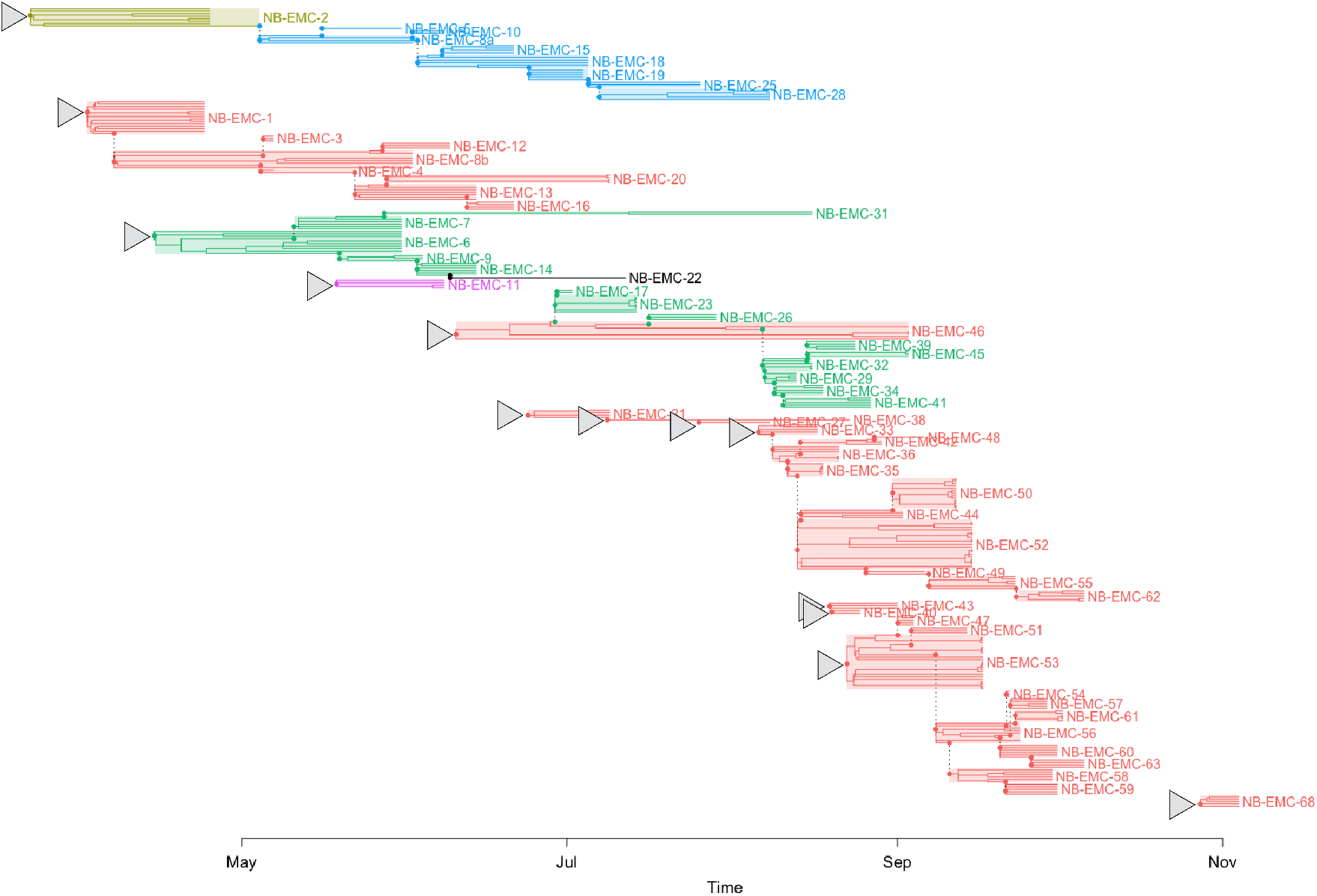
Maximum parent credibility transmission tree with with-host phylogenetic trees for SARS-CoV-2 outbreak in mink farms. The farms are colored according to the clusters found by Lu et al. (2021): cluster A: red; cluster B; yellow, cluster C: green; cluster D: blue, cluster E: purple, cluster unknown: black. Cluster A is divided into 5 smaller clusters, with cluster A1 introduced in NB-EMC-1 and cluster A2 introduced in NB-EMC-46.

**Figure S6:**
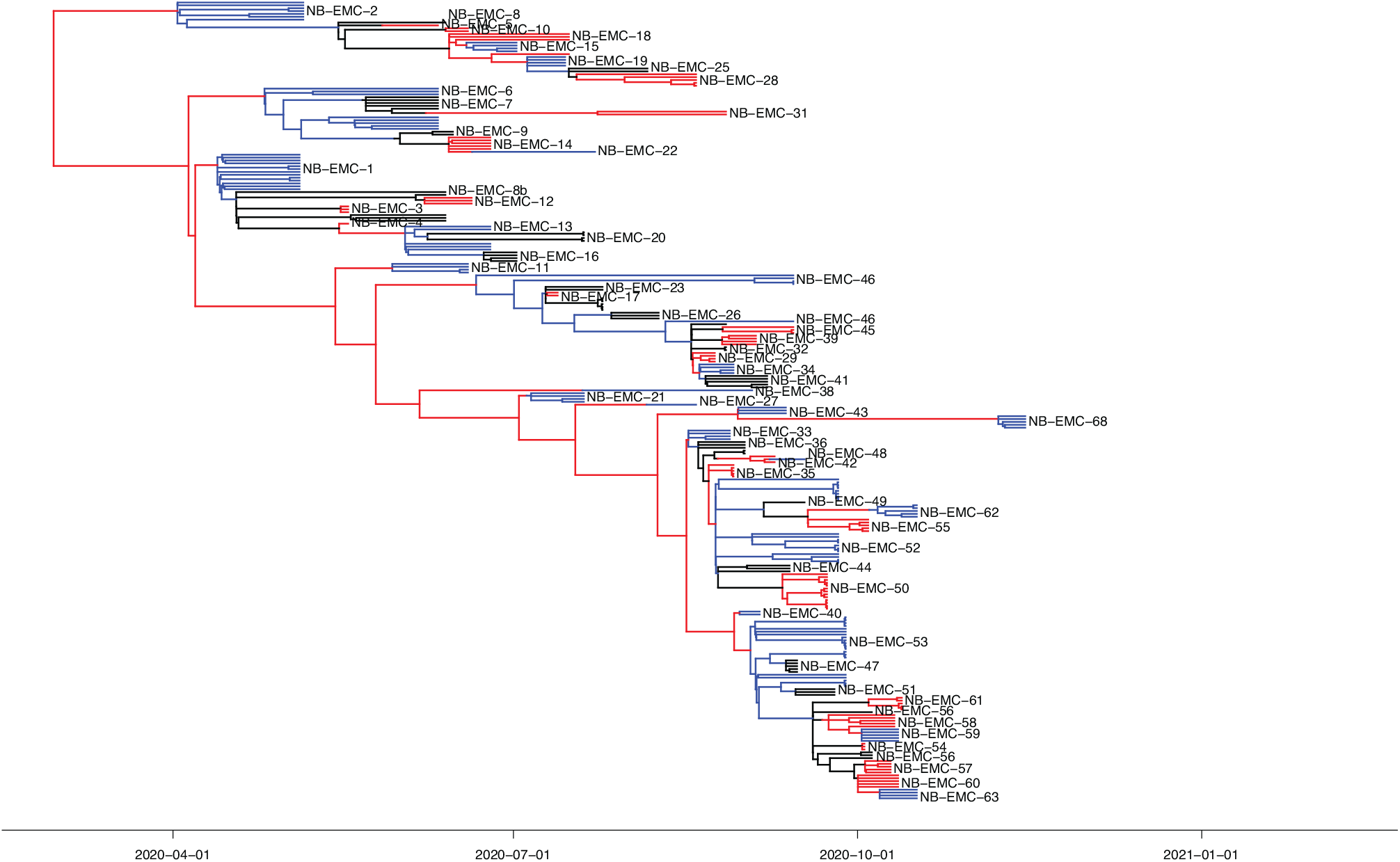
Maximum parent credibility phylogenetic tree for SARS-CoV-2 outbreak in mink farms. The history host is shown as the most-left red line, and the hosts are given in alternating colors. The black boxes represent the clusters in the transmission tree, with the lowest box the assumed bigger cluster with index case NB-EMC-46.

**Figure S7:**
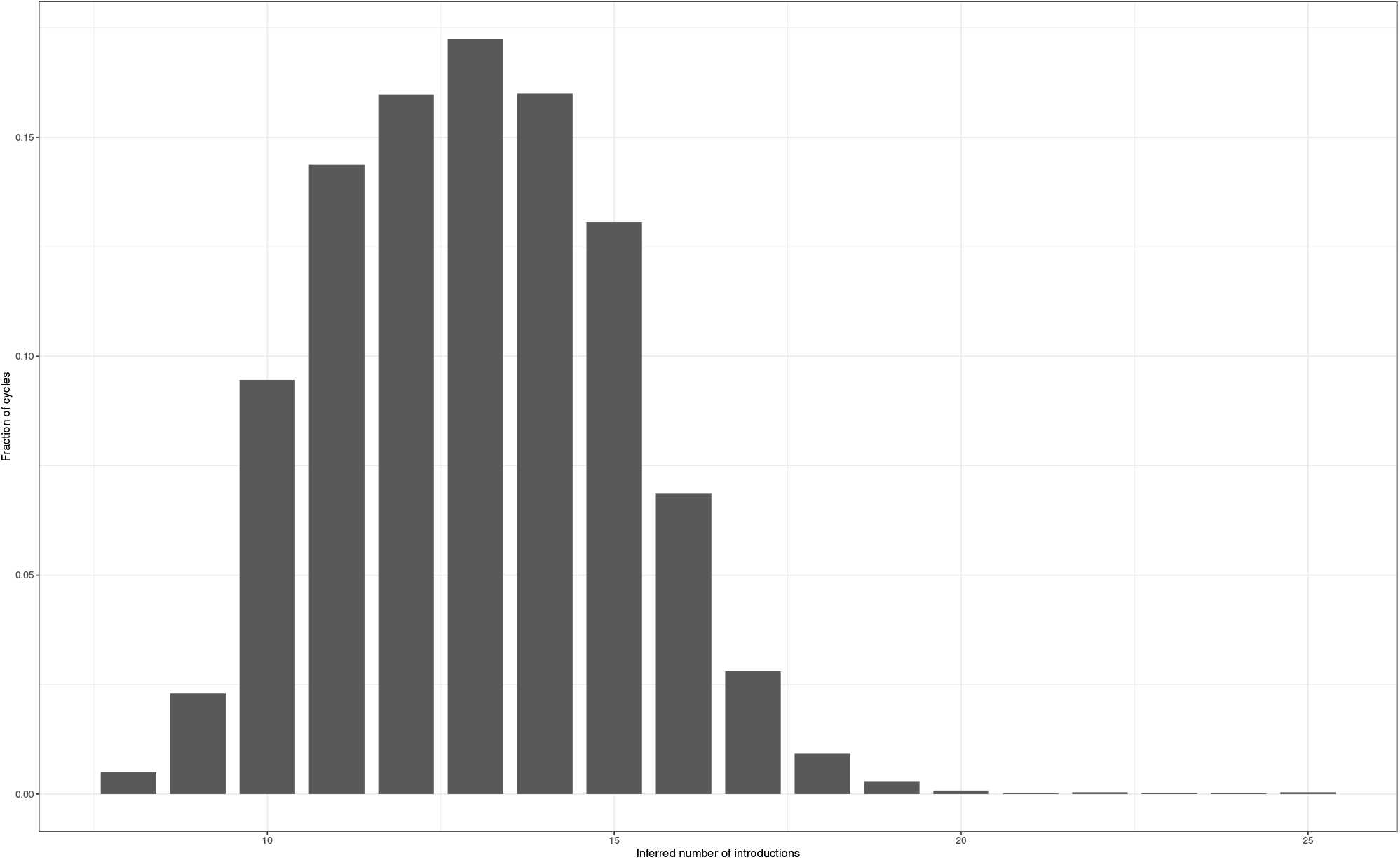
Histogram of number of introductions for the mink farms.

**Figure S8:**
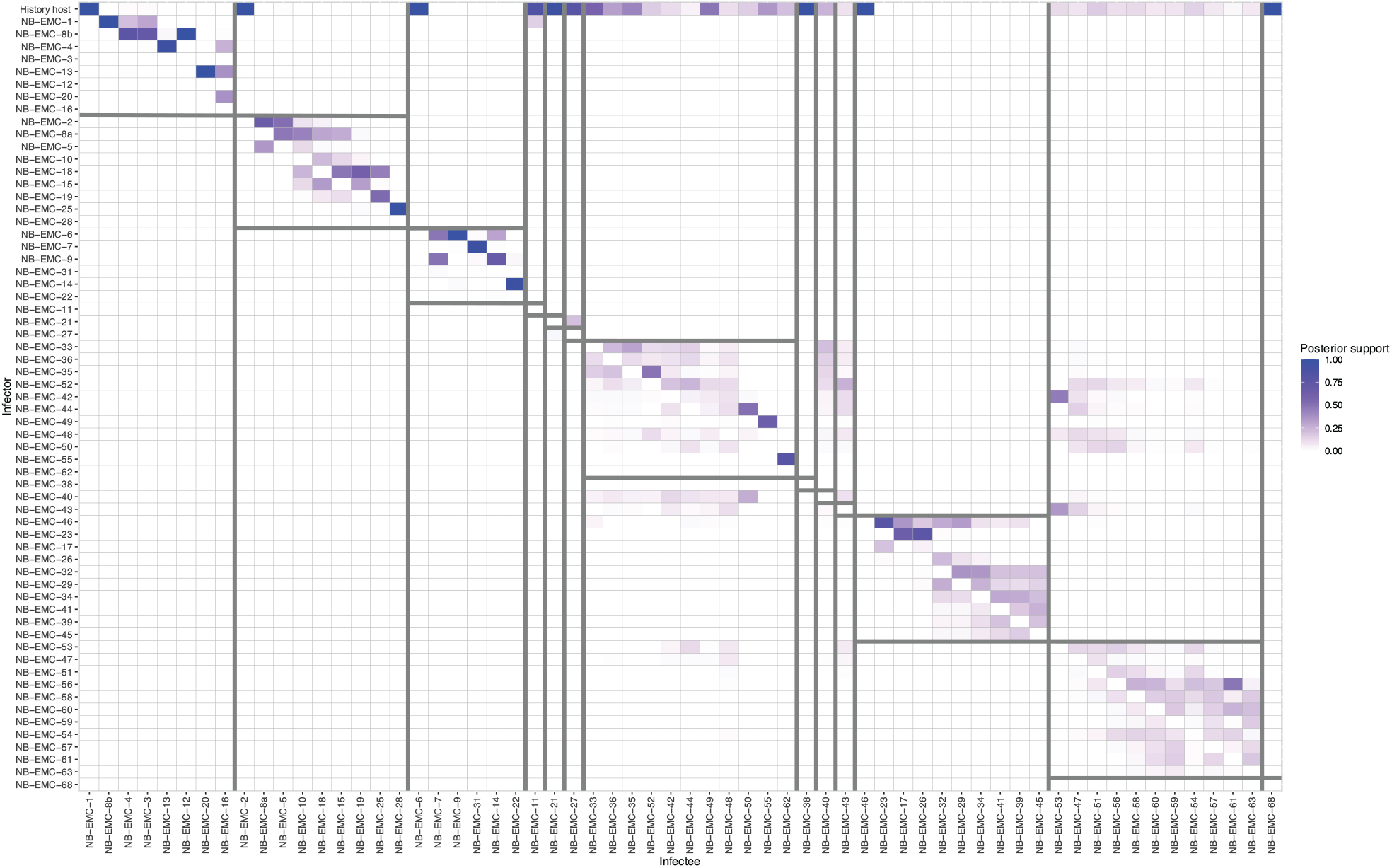
Posterior support of infectors of all hosts. There is a high certainty of the index cases (infectees with the history host as infector) in the beginning of the outbreak. Transmission clusters with index cases NB-EMC-33 and NB-EMC-53 show more variation of the infectors, even outside their transmission cluster. Posterior support is shown from 0 (white) to 1 (blue). Hosts are ordered by transmission cluster and infection time. The grey bars show the transmission clusters.

**Table S1:** Comparison between MCMC and MC^3^. Differences between median posterior log-likelihood and the log-likelihood of the simulated outbreak. Results are the means from analyses of 25 outbreaks for each setting, of the 10,001st to 35,000th MCMC cycle of each outbreak analysis.

| Simulated numbers of introductions | Method | Log-likelihood |
| --- | --- | --- |
| 4*1 | MCMC random | 68.3 |
|  | p(MC<sup>3</sup>) random | 70.1 |
|  | MCMC NJ | 67.9 |
|  | p(MC<sup>3</sup>) NJ | 70.1 |
| 4*5 | MCMC random | 27.3 |
|  | p(MC<sup>3</sup>) random | 27.0 |
|  | MCMC NJ | 27.5 |
|  | p(MC<sup>3</sup>) NJ | 27.0 |
| 4*10 | MCMC random | **9.78** |
|  | p(MC<sup>3</sup>) random | 20.0 |
|  | MCMC NJ | 17.0 |
|  | p(MC<sup>3</sup>) NJ | 20.2 |
| 4*15 | MCMC random | **-2.04** |
|  | p(MC<sup>3</sup>) random | 14.5 |
|  | MCMC NJ | 9.49 |
|  | p(MC<sup>3</sup>) NJ | 14.5 |

**Table S2:** Inferring multiple introductions with varying prior information: no information, informative priors, and fixed parameters. 25 outbreaks of size 20 are simulated with 5 introductions for each set of priors. The results of the model parameters are mean differences between mean estimates and the simulated value.

|  | No information <sup>c</sup> | Informative priors <sup>b</sup> | Fixed parameters <sup>a</sup> |
| --- | --- | --- | --- |
| **Mean difference between estimations and simulated value** |  |  |  |
| Introductions | 0.16 | 0.12 | 0.41 |
| $\mu$ | $3.28 \cdot 10^{-5}$ | $4.59 \cdot 10^{-6}$ | 0 |
| $m_G$ | 0.22 | 0.05 | 0 |
| $m_S$ | 0.40 | 0.02 | 0 |
| $r$ | 0.08 | 0.10 | 0 |
| $r_{\text{history}}$ | 15.1 | 4.95 | 0 |
| **Tree inference** |  |  |  |
| True infectors with highest support | 15/20 | 15/20 | 15.7/20 |
| True infectors in 95% CI | 19.8/20 | 20/20 | 19.5/20 |

<sup>a</sup> $\mu_G = 1, \sigma_G = \infty, \mu_S = 1, \sigma_S = \infty, \mu_\mu = 0, \sigma_\mu = 100$ <sup>b</sup> $\mu_G = 1, \sigma_G = 0.1, \mu_S = 1, \sigma_S = 0.1, \mu_\mu = 10^{-4}, \sigma_\mu = 5 \cdot 10^{-5}$ <sup>c</sup> $m_G, m_S, r = 1, r_{\text{history}} = 50, \mu = 10^{-4}$

**Table S3:**
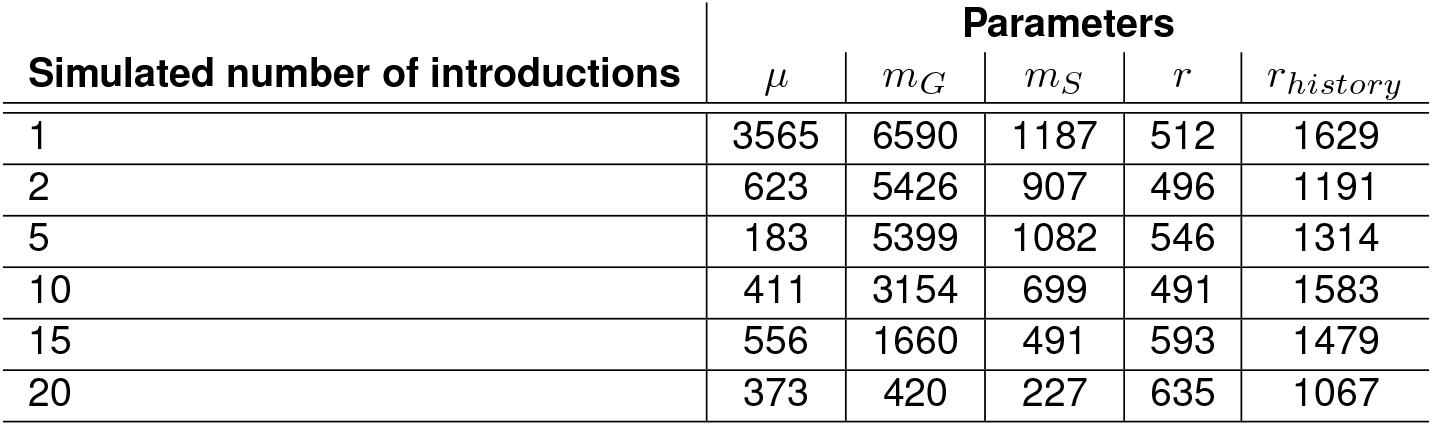
Effective Sample Sizes of the model parameters calculated for a various number of introductions. Results are the mean of 75 chains, i.e. 3 coalescent rates per number of introductions and 25 outbreaks per parameter set.

**Table S4:**
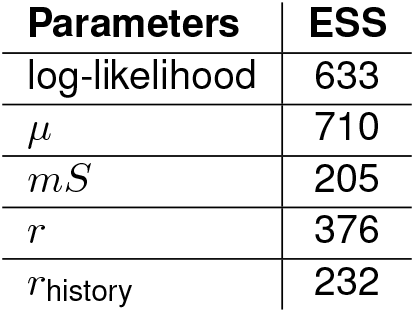
Effective Sample Sizes (ESS) of the model parameters for analyzing a SARS-CoV-2 outbreak in mink farms in the Netherlands.

## Notes

### Competing Interest Statement

The authors have declared no competing interest.

## References

1. Altekar, G., Dwarkadas, S., Huelsenbeck, J. P., and Ronquist, F. Parallel Metropolis coupled Markov chain Monte Carlo for Bayesian phylogenetic inference. Bioinformatics, 20(3):407–415, 2004. doi: 10.1093/bioinformatics/btg427.

2. Amicone, M., Borges, V., Alves, M. J., Isidro, J., Zé-Zé, L., Duarte, S., Vieira, L., Guiomar, R., Gomes, J. P., and Gordo, I. Mutation rate of SARS-CoV-2 and emergence of mutators during experimental evolution. Evolution, Medicine, and Public Health, 10(1):142–155, 1 2022. doi: 10.1093/emph/eoac010.

3. Cauchemez, S. and Ferguson, N. M. Methods to infer transmission risk factors in complex outbreak data. Journal of the Royal Society, Interface, 9(68):456–69, 3 2012. doi: 10.1098/rsif.2011.0379.

4. Cauchemez, S., Boelle, P.-Y., Donnelly, C. A., Ferguson, N. M., Thomas, G., Leung, G. M., Hedley, A. J., Anderson, R. M., and Valleron, A.-J. Real-time estimates in early detection of SARS. Emerging infectious diseases, 12(1):110–3, 1 2006. doi: 10.3201/eid1201.050593.

5. Didelot, X., Gardy, J., and Colijn, C. Bayesian inference of infectious disease transmission from whole-genome sequence data. Molecular Biology and Evolution, 2014. doi: 10.1093/molbev/msu121.

6. Didelot, X., Fraser, C., Gardy, J., Colijn, C., and Malik, H. Genomic infectious disease epidemiology in partially sampled and ongoing outbreaks. Molecular Biology and Evolution, 34(4):997–1007, 4 2017. doi: 10.1093/molbev/msw275.

7. Edgar, R. C. MUSCLE: multiple sequence alignment with high accuracy and high throughput. Nucleic Acids Research, 32 (5):1792–1797, 3 2004. doi: 10.1093/nar/gkh340.

8. Felsenstein, J. Evolutionary trees from DNA sequences: A maximum likelihood approach. Journal of Molecular Evolution, 17(6):368–376, 11 1981. doi: 10.1007/BF01734359.

9. Fraser, C., Donnelly, C. A., Cauchemez, S., Hanage, W. P., Van Kerkhove, M. D., Hollingsworth, T. D., Griffin, J., Baggaley, R. F., Jenkins, H. E., Lyons, E. J., Jombart, T., Hinsley, W. R., Grassly, N. C., Balloux, F., Ghani, A. C., Ferguson, N. M., Rambaut, A., Pybus, O. G., Lopez-Gatell, H., Alpuche-Aranda, C. M., Chapela, I. B., Zavala, E. P., Guevara, D. M. E., Checchi, F., Garcia, E., Hugonnet, S., and Roth, C. Pandemic Potential of a Strain of Influenza A (H1N1): Early Findings. Science, 324(5934):1557–1561, 6 2009. doi: 10.1126/science.1176062.

10. Hall, M., Woolhouse, M., and Rambaut, A. Epidemic Reconstruction in a Phylogenetics Framework: Transmission Trees as Partitions of the Node Set. PLoS Computational Biology, 11(12):1–36, 2015. doi: 10.1371/journal.pcbi.1004613.

11. Hammer, A. S., Quaade, M. L., Rasmussen, T. B., Fonager, J., Rasmussen, M., Mundbjerg, K., Lohse, L., Strandbygaard, B., Jørgensen, C. S., Alfaro-Núñez, A., Rosenstierne, M. W., Boklund, A., Halasa, T., Fomsgaard, A., Belsham, G. J., and Bøtner, A. SARS-CoV-2 Transmission between Mink (Neovison vison) and Humans, Denmark. Emerging Infectious Diseases, 27(2):547–551, 2 2021. doi: 10.3201/eid2702.203794.

12. Harris, S. R., Feil, E. J., Holden, M. T. G., Quail, M. A., Nickerson, E. K., Chantratita, N., Gardete, S., Tavares, A., Day, N., Lindsay, J. A., Edgeworth, J. D., de Lencastre, H., Parkhill, J., Peacock, S. J., and Bentley, S. D. Evolution of MRSA During Hospital Transmission and Intercontinental Spread. Science, 327(5964):469–474, 1 2010. doi: 10.1126/science.1182395.

13. Haydon, D. T., Chase-Topping, M., Shaw, D. J., Matthews, L., Friar, J. K., Wilesmith, J., and Woolhouse, M. E. J. The construction and analysis of epidemic trees with reference to the 2001 UK foot-and-mouth outbreak. Proceedings. Biological sciences, 270(1511):121–7, 1 2003. doi: 10.1098/rspb.2002.2191.

14. Jombart, T., Cori, A., Didelot, X., Cauchemez, S., Fraser, C., and Ferguson, N. Bayesian Reconstruction of Disease Outbreaks by Combining Epidemiologic and Genomic Data. PLoS Computational Biology, 10(1), 2014. doi: 10.1371/journal.pcbi.1003457.

15. Kenah, E., Britton, T., Halloran, M. E., and Longini, I. M. Molecular Infectious Disease Epidemiology: Survival Analysis and Algorithms Linking Phylogenies to Transmission Trees. PLoS computational biology, 12(4):e1004869, 4 2016. doi: 10.1371/journal.pcbi.1004869.

16. Kenah, E. Semiparametric Relative-risk Regression for Infectious Disease Transmission Data. Journal of the American Statistical Association, 110(509):313–325, 3 2015. doi: 10.1080/01621459.2014.896807.

17. Kerfua, S. D., Shirima, G., Kusiluka, L., Ayebazibwe, C., Mwebe, R., Cleaveland, S., and Haydon, D. Spatial and temporal distribution of foot-and-mouth disease in four districts situated along the Uganda-Tanzania border: Implications for cross-border efforts in disease control. The Onderstepoort journal of veterinary research, 85(1):e1–e8, 8 2018. doi: 10.4102/ojvr.v85i1.1528.

18. Klinkenberg, D., Backer, J. A., Didelot, X., Colijn, C., and Wallinga, J. Simultaneous inference of phylogenetic and transmission trees in infectious disease outbreaks. PLoS Computational Biology, 13(5), 5 2017. doi: 10.1371/journal.pcbi.1005495.

19. Lu, L., Sikkema, R. S., Velkers, F. C., Nieuwenhuijse, D. F., Fischer, E. A., Meijer, P. A., Bouwmeester-Vincken, N., Rietveld, A., Wegdam-Blans, M. C., Tolsma, P., Koppelman, M., Smit, L. A., Hakze-van der Honing, R. W., van der Poel, W. H., van der Spek, A. N., Spierenburg, M. A., Molenaar, R. J., Rond, J. d., Augustijn, M., Woolhouse, M., Stegeman, J. A., Lycett, S., Oude Munnink, B. B., and Koopmans, M. P. Adaptation, spread and transmission of SARS-CoV-2 in farmed minks and associated humans in the Netherlands. Nature Communications, 12(1), 12 2021. doi: 10.1038/s41467-021-27096-9.

20. Morelli, M. J., Thébaud, G., Chadœuf, J., King, D. P., Haydon, D. T., and Soubeyrand, S. A Bayesian Inference Framework to Reconstruct Transmission Trees Using Epidemiological and Genetic Data. PLoS Computational Biology, 8(11):e1002768, 11 2012. doi: 10.1371/journal.pcbi.1002768.

21. Munnink, B. B., Sikkema, R. S., Nieuwenhuijse, D. F., Molenaar, R. J., Munger, E., Molenkamp, R., Van Der Spek, A., Tolsma, P., Rietveld, A., Brouwer, M., Bouwmeester-Vincken, N., Harders, F., Der Honing, R. H. V., Wegdam-Blans, M. C., Bouwstra, R. J., GeurtsvanKessel, C., Van Der Eijk, A. A., Velkers, F. C., Smit, L. A., Stegeman, A., Van Der Poel, W. H., and Koopmans, M. P. Transmission of SARS-CoV-2 on mink farms between humans and mink and back to humans. Science, 371(6525):172–177, 1 2021. doi: 10.1126/science.abe5901.

22. Mutreja, A., Kim, D. W., Thomson, N. R., Connor, T. R., Lee, J. H., Kariuki, S., Croucher, N. J., Choi, S. Y., Harris, S. R., Lebens, M., Niyogi, S. K., Kim, E. J., Ramamurthy, T., Chun, J., Wood, J. L. N., Clemens, J. D., Czerkinsky, C., Nair, G. B., Holmgren, J., Parkhill, J., and Dougan, G. Evidence for several waves of global transmission in the seventh cholera pandemic. Nature, 477(7365):462–465, 9 2011. doi: 10.1038/nature10392.

23. Numminen, E., Chewapreecha, C., Sirén, J., Turner, C., Turner, P., Bentley, S. D., and Corander, J. Two-phase importance sampling for inference about transmission trees. Proceedings of the Royal Society B: Biological Sciences, 281(1794), 2014. doi: 10.1098/rspb.2014.1324.

24. O’Toole, Scher, E., Underwood, A., Jackson, B., Hill, V., McCrone, J. T., Colquhoun, R., Ruis, C., Abu-Dahab, K., Taylor, B., Yeats, C., Du Plessis, L., Maloney, D., Medd, N., Attwood, S. W., Aanensen, D. M., Holmes, E. C., Pybus, O. G., and Rambaut, A. Assignment of Epidemiological Lineages in an Emerging Pandemic Using the Pangolin Tool. Virus Evolution, 7 2021. doi: 10.1093/ve/veab064.

25. Pham, T. M., Kretzschmar, M., Bertrand, X., and Bootsma, M. Tracking Pseudomonas aeruginosa transmissions due to environmental contamination after discharge in ICUs using mathematical models. PLOS Computational Biology, 15(8): e1006697, 8 2019. doi: 10.1371/journal.pcbi.1006697.

26. R Core Team. R: A Language and Environment for Statistical Computing, 2022. URL https://www.r-project.org/.

27. Ruan, Y., Wei, C. L., Ling, A. E., Vega, V. B., Thoreau, H., Se Thoe, S. Y., Chia, J.-M., Ng, P., Chiu, K. P., Lim, L., Zhang, T., Chan, K. P., Lin Ean, L. O., Ng, M. L., Leo, S. Y., Ng, L. F., Ren, E. C., Stanton, L. W., Long, P. M., and Liu, E. T. Comparative full-length genome sequence analysis of 14 SARS coronavirus isolates and common mutations associated with putative origins of infection. The Lancet, 361(9371):1779–1785, 5 2003. doi: 10.1016/S0140-6736(03)13414-9.

28. Si, Y., de Boer, W. F., and Gong, P. Different environmental drivers of highly pathogenic avian influenza H5N1 outbreaks in poultry and wild birds. PloS one, 8(1):e53362, 2013. doi: 10.1371/journal.pone.0053362.

29. Worby, C. J., O’Neill, P. D., Kypraios, T., Robotham, J. V., De Angelis, D., Cartwright, E. J., Peacock, S. J., and Cooper, B. S. Reconstructing transmission trees for communicable diseases using densely sampled genetic data. Annals of Applied Statistics, 10(1):395–417, 3 2016. doi: 10.1214/15-AOAS898.

30. Ypma, R. J., van Ballegooijen, W. M., and Wallinga, J. Relating phylogenetic trees to transmission trees of infectious disease outbreaks. Genetics, 195(3):1055–1062, 2013. doi: 10.1534/genetics.113.154856.

31. Zhao, S., Tang, B., Musa, S. S., Ma, S., Zhang, J., Zeng, M., Yun, Q., Guo, W., Zheng, Y., Yang, Z., Peng, Z., Chong, M. K., Javanbakht, M., He, D., and Wang, M. H. Estimating the generation interval and inferring the latent period of COVID-19 from the contact tracing data. Epidemics, 36, 2021. doi: 10.1016/j.epidem.2021.100482.

